# Decoding auditory working memory load from EEG alpha oscillations

**DOI:** 10.1101/2025.02.14.638227

**Authors:** Yichen Yuan, Surya Gayet, Derk Christiaan Wisman, Stefan van der Stigchel, Nathan van der Stoep

## Abstract

Working memory (WM) enables temporary retention of task-relevant information for imminent use. Increases in visual WM load are accompanied by elevated contralateral delay activity (CDA), and EEG alpha-band power. While most WM research focuses on the visual domain, it remains unknown whether similar EEG responses also reflect WM load in the auditory domain. Using EEG, we set out to establish such neural markers of auditory WM load. Participants memorized the pitches of 1 to 4 pure tones presented to one ear, with 1 to 4 identical distractor tones presented to the other ear. Behaviorally, auditory WM capacity plateaued between set-sizes two and three. Unlike for visual WM, auditory WM load was not reflected in lateralized EEG responses. This shows that the CDA is a vision-specific rather than domain-general neural marker of WM load. Applying multivariate pattern analyses on the delay activity revealed that auditory WM load is reflected in patterns of alpha-band oscillations. Surprisingly, a temporal generalization analysis revealed that the alpha patterns reflecting specific load conditions changed throughout the maintenance period (despite load being inherently constant), revealing dynamic coding of auditory WM load.

## 1 Introduction

Working memory (WM) is a limited-capacity system that allows us to temporarily hold several representations accessible in service of other mental tasks (Cowan, 1998). Commonly, not all task-relevant information is available in one location or at a single moment, which requires us to temporarily retain information in mind. For instance, we may memorize a phone number for subsequent dialing or remembering a list of ingredients while grocery shopping. WM functions as our central information storage-and-processing structure (Cowan et al., 2005), which enables us to interact with the world over space and time.

So far, Baddeley and Hitch (1974) proposed probably the most influential WM model, which comprises three components: (1) a visuo-spatial sketchpad for storage of visual information; (2) a phonological loop for storage of auditory information; (3) a central executive that regulates the content of the active portion of WM. Years later, Baddeley (2000) added a fourth component, the episodic buffer, that binds features from different sources together into multidimensional objects. According to Baddeley’s model, the storage of WM is modality-specific, with a visuo-spatial sketchpad for visual information and a phonological loop for auditory information. Indeed, previous studies have provided evidence that visual and auditory WM may rely on at least partly distinct structures that produce dissociable neural responses (Lefebvre et al., 2013; Pratt et al., 1989; also see Scimeca et al., 2018 for sensory recruitment hypothesis).

Given the (at least partly) modality-specific nature of WM, it is reasonable to use modality-specific stimuli to study visual WM and auditory WM separately. Yet, most WM studies to date focused on the visual modality, for instance the color, orientation, spatial location, etc. of visual items (Carlisle et al., 2011; Diamantopoulou et al., 2011; Harrison & Tong, 2009; Li & Saiki, 2015; Luria & Vogel, 2011; Schurgin, 2018). Correspondingly, studies have found evidence for the existence of a ‘magic number four’, the observation that visual working memory capacity is severely limited to an average of four items in young adults (Cowan, 2010). By recording electroencephalograms (EEG), Luck and Vogel (1997) identified a neural marker of visual WM load and named it Contralateral Delay Activity (CDA). Specifically, they presented a symmetric visual array in which the left and right side differed in features and instructed the participants to memorize the objects in only one hemifield. In this bilateral change-detection paradigm, information presented on both sides was perceptually processed, but information presented in only one hemifield was subsequently retained in WM. Interhemispheric difference waves were calculated for each set-size condition by subtracting the ipsilateral activity from the contralateral activity. By doing so, any nonspecific, bilateral Event-Related Potentials (ERPs) were removed. This approach thus allows to isolate the neural activity specifically related to encoding and maintenance of the memorized objects. A large negative-going voltage over the contralateral hemisphere (relative to memorized hemifield) was observed, located primarily over posterior parietal and lateral occipital electrodes. Moreover, the amplitude of the CDA varied with visual WM load, as it increased between set-sizes of one, two and three items, and then leveling off at the group’s average capacity limit of about three items (Vogel & Machizawa, 2004). Interestingly, the existence of lateralized responses to set-size (such as the CDA), imply that visual WM is inherently spatially organized.

Another neural marker of visual WM has been observed in the frequency domain, particularly in the alpha band, although studies have reported contradictory results regarding it’s modulation direction. For instance, Jensen et al. (2002) have shown that the power of oscillations in the alpha-band (8-12 Hz) over the posterior and central EEG channels track visual memory load during the maintenance period. Such increases in alpha band power during WM maintenance are widely accepted as reflecting functional disengagement or inhibition of task-irrelevant visual inputs to protect the task-relevant information maintained in WM (Bonnefond & Jensen, 2012; Roux & Uhlhaas, 2014; Tuladhar et al., 2007; Wianda & Ross, 2019). In contrast, a number of studies have reported attenuated alpha oscillations in sensory areas that are mnemonically relevant, suggesting that alpha decreases may support the recruitment of these areas for perceptual WM retention (Fukuda et al., 2015; van Ede, 2018). Taken together, several neural markers of WM load have been established in the visual domain, some of which are lateralized (capitalizing on the spatial organization of visual WM) and some of which are not.

Surprisingly, much less is known about auditory WM compared to visual WM. In the current study, we define auditory WM as the maintenance of acoustic properties of sound stimuli, such as the pitch, duration, timbre, and amplitude (following Lefebvre et al., 2013). It should be noted that the maintenance of verbal information is not necessarily the same as auditory WM. Studies have found that maintenance of verbal and of purely acoustic material share relatively few characteristics (Deutsch, 1970; Williamson et al., 2010). In the studies focusing on the maintenance of acoustic properties, the auditory WM capacity found in tasks using pure tones was around 2 to 3 (Alunni-Menichini et al., 2014; Li et al., 2013; Prosser, 1995). Lefebvre et al. (2013) set out to identify the neural marker of auditory short-term memory. Contrasting the univariate EEG response during the maintenance period between the memory task and a control task revealed a sustained negative-going voltage over the central- frontal electrodes that scaled with auditory WM load. This negativity was identified as a neural marker of auditory WM and named as the sustained anterior negativity (SAN). In a follow-up study, Alunni-Menichini et al. (2014) showed that the amplitude of SAN increased in negativity from 2 to 4 items and then levelled off from 4 to 8 items. Thus, they established the SAN as an index of brain activity specifically reflecting the amount of information (load) maintained in auditory WM. In these studies investigating auditory WM, however, stimulus presentation was not lateralized, unlike typical visual WM studies measuring lateralized responses (CDA). Thus, it remains unknown whether lateralized responses that hinge on the inherently spatial organization of WM (akin to the CDA for visual WM) can also be observed for auditory WM load. On the one hand, it is reasonable to assume that the visual CDA reflects an enhanced representation of the contralateral spatial location where the memorized items are stored (e.g. Klaver et al., 1999, Talsma et al., 2001), suggesting a modality-specific neural marker for visual WM alone. On the other hand, if the CDA reflects a more abstract form of working memory representation, the lateralized CDA-like responses should also be observed in auditory working memory task.

Other studies, focusing on the frequency domain of the EEG response, have found that alpha oscillations (8-12 Hz) are also a sensitive marker for auditory WM load (Kaiser et al., 2007; Leiberg et al., 2006; Luo et al., 2005; Van Dijk et al., 2010). For instance, Leiberg et al. (2006) found monotonic increases in spectral amplitude as a function of memory load for the alpha band over right frontal sensors during the delay period. Similarly, Van Dijk et al. (2010) also found a left-lateralized (left temporal regions) increase in 5-12 Hz during the maintenance of pitches, compared to a non-memory control task. Alpha-band oscillations can therefore be considered to reflect the maintenance of auditory information. Earlier work has interpreted alpha-band oscillatory responses to either reflect the top-down control of a WM representations (Leiberg et al., 2006) or the inhibition of task-irrelevant neural processes (Van Dijk et al., 2010). Either way, alpha-band oscillatory responses are closely related to maintenance of auditory information (for a review, see Wilsch & Obleser, 2016), and may therefore track auditory WM load.

Recently, multivariate pattern analysis (MVPA) has been advocated in the study of higher order brain states, as it is more sensitive to pick up on complex scalp patterns of neural activity, compared to univariate methods (Kikumoto & Mayr, 2018; Peelen & Downing, 2023). While most WM decoding studies have focused on visual representations, less is known about the maintenance of auditory stimulus features. A handful of multivariate fMRI studies have shown that auditory information (e.g., frequency, location, etc.) can be decoded in auditory cortex, inferior frontal cortex, precentral cortex, and superior parietal lobule (for a review, see Kaiser, 2015). Moreover, we are not aware of any study using multivariate analyses to investigate executive properties (such as load) of or auditory WM. Thus, in the present study we set out to uncover the spatio-temporal patterns of EEG activity relating to auditory WM load.

Considering the large knowledge gap between what we know about visual WM and auditory WM, it seems imperative to study auditory WM via various methods (especially multivariate methods) used in visual WM literatures. Moreover, previous studies into auditory WM have only used binaural stimuli, where potential lateralized (i.e., contralateral-minus-ipsilateral) responses akin to the CDA would remain unnoticed. It therefore remains unknown whether the CDA is a modality-specific neural marker for visual WM or whether it is a supra-modal neural marker that also reflects auditory WM load. In the current study, we aimed to identify load-dependent neural markers of auditory WM by recording EEG signals. Specifically, we asked two research questions, namely, (1) whether auditory WM load is reflected in lateralized neural responses, akin to the CDA for visual WM load, and (2) whether we could identify non-lateralized load-dependent neural markers of auditory WM, using multivariate pattern analyses.

To address these questions, we conducted an auditory delayed change-detection task (e.g., Rouder et al., 2011) while recording EEG. On each trial, participants memorized a tone sequence comprising 1, 2, 3, or 4 pure tones differing in pitch, thus yielding four set-sizes. Importantly, this auditory change-detection task was combined with a dichotic listening task (e.g., Hugdahl, 2011), whereby the to-be-memorized tone sequences were presented to one ear, while a to-be-ignored distractor sequence was presented to the other ear. Participants were instructed to memorize the tones presented to one ear, while ignoring the tones presented to the other ear. After a delay period, a probe sound was presented to the attended ear, with a distractor presented to the un-attended ear. Participants were required to indicate whether the probe was present or absent in the memory tone sequence. This dichotic listening approach allowed us to isolate load-dependent and hemisphere-specific EEG responses during the delay period of the auditory WM task. By doing so, we could compute the contralateral-minus-ipsilateral response differences, and compare these between set-size conditions to identify potential lateralized load-depended neural markers of auditory WM (akin to the CDA).

We analyzed the EEG data using both univariate and multivariate methods (EEG and decoding), investigating lateralized and non-lateralized responses, in the time and frequency domain. EEG responses are retrieved from the maintenance period, when participants held varying numbers of pitches (i.e., set-sizes) in auditory WM. For behavioral performance, we expected that the estimation of WM capacity should increase with set-size, levelling off at the WM capacity limit. For the univariate EEG results, we focused on lateralized responses, namely the CDA responses and lateralized alpha oscillations, based on previous visual WM studies (Jensen et al., 2002; Vogel & Machizawa, 2004). If auditory WM is also reflected in lateralized responses, the contralateral-minus-ipsilateral univariate responses and alpha-band power should scale with auditory WM set-size (until it levels off at the WM capacity limit). For multivariate pattern analysis, we focused on decoding scalp patterns of alpha-band (8-12 Hz) power, based on previous findings that alpha was closely related to auditory WM (Leiberg et al., 2006; Wilsch & Obleser, 2016). If the patterns of alpha oscillation during the maintenance period indeed track auditory WM load, the alpha oscillation patterns should be distinguishable between load conditions up to the group-level capacity limitations, but fail to differentiate among conditions that exceeded this limit, as reflected in the auditory WM capacity calculated by the behavioral data.

To preface the results, we found that auditory WM load was not reflected in lateralized responses, neither in the time nor in the frequency domain. This implies that the CDA is vision-specific rather than domain-general marker of WM load. Our decoding results showed that patterns of alpha-band oscillations during the maintenance period reflected auditory WM load. Moreover, the alpha patterns associated with specific load conditions were changing throughout the maintenance period, which suggests that auditory WM load is reflected in dynamic – rather than static – neural population codes. Mirroring the behavioral data that the auditory WM capacity is around 2 tones, these load-specific EEG responses allowed to distinguish between load conditions within, but not beyond, the group-level capacity limitations. We thus consider scalp patterns of alpha band power as a novel neural marker for auditory WM load.

## 2 Materials and Methods

### 2.1 Participants

To determine the required sample size for a main effect of set-size in our within-participants design, a sample size estimation was performed in G*power software (Faul et al., 2009). This suggested that at least 19 participants were required for 85% power to observe a medium effect size (Cohen’s *f* = .3) with a repeated measure ANOVA (a = 0.05). The medium effect size was inferred based on (1) a general benchmark that is often used when effect sizes are a priori unknown, (2) the neural effect sizes inferred from Vogel & Machizawa (2004), and (3) one of our piloting behavioral experiment with the same design but having set-size 1, 2, 3, 4, 5, and 6. A total of 27 participants were tested (18 participants reported their gender as female, 9 as male; mean age = 22.41 years, *SD* = 1.91, range = 19 to 25 years). We stopped testing after enough participants (>19) met our inclusion criteria for the EEG analyses (for details, see the EEG recording section below). Six participants were excluded due to excessive EEG artifacts (> 25%). Thus, EEG data of 21 participants were analyzed. All participants reported normal or corrected-to-normal vision and normal hearing (as tested with a pre-experiment audiogram). They signed informed consent and received money or course credits for their participation. The study protocols were approved by the faculty ethics committee (FETC) of Utrecht University (number 18-048 van der Stoep).

### 2.2 Apparatus and stimuli

The experiments were conducted in a dimly lit lab and controlled using Matlab 2018a. Participants were seated with their head positioned in a chin rest, to keep their viewing distance at a fixed 60 cm in front of a 27-inch monitor. The auditory stimuli were played through 3M™ E-A-RTONE™ Insert Earphone 3A (10 Ohm).

The auditory stimuli consisted of 25 pure tones with different frequencies and white noises, each lasting for 200 ms. All sound were generated in Matlab 2018a, sampled at 96 kHz. The frequencies of the 25 target pure tones ranged from 125 Hz to 8.1 kHz with a 19% increase in-between tones (125, 149, 177, 211, 251, 298, 355, 422, 503, 598, 712, 847, 1008, 1200, 1428, 1699, 2021, 2406, 2863, 3407, 4054, 4824, 5740, 6831, 8129 Hz), which should be distinguishable for naive listeners (Ahissar et al., 2006). The lowest (125 Hz) and highest (8129 Hz) frequency from this set of pure tones were used as distractors, while the remaining 23 pure tones were used as the set of target stimuli. Linear ramps of 25 ms were applied both to the beginning and the end of tones to prevent auditory pop artifacts. Participants were able to slightly adjust the volume to a subjectively comfortable level at the start of the WM tasks. Sounds were on average presented at 60 dB(A).

### 2.3 Procedure

#### Pitch discrimination task

To make sure that participants could distinguish different pure tones, a behavioral pitch discrimination task was conducted before the main auditory WM task. In this pitch discrimination task, each of the 25 pre-generated pure tones was paired with itself, yielding 25 same-frequency trials. In addition, 23 pure tones (except for the lowest and highest ones) were paired with their one-step higher or lower neighbors, yielding 46 different-frequency trials. Finally, the lowest tone was paired with its one-step higher neighbor, and the highest tone with its one-step lower neighbor, yielding 2 additional different-frequency trials. This resulted in a total of 73 trials (pairs), consisting of 25 same-frequency and 48 different-frequency trials. On each trial, one pair of tones was presented bilaterally and sequentially. Participants were required to indicate whether the two tones were the same or different. All pairs were presented once. Accuracies in the task for all participants were higher than 85% (M = 97.2%, *SD* = 2.86%, range = 89% to 100%), indicating that participants could distinguish the different tones; therefore, no participants were excluded based on this criterion.

#### Auditory WM task

After the pitch discrimination task, participants performed the auditory WM task during EEG recording. In this WM task, a 4 (Set-size: 1, 2, 3, vs. 4) × 2 (Attended side: Left vs. Right) within-participants design was adopted. At the beginning of each trial, participants were always given instructions to remind them of which side to attend to. They were required to memorize the pitches of a sound sequence presented on the attended side, while ignoring the sound sequence presented on the unattended side. The manipulation of the to-be-attended side was blocked, while set-size varied randomly from trial to trial. The order of blocks was counterbalanced across participants.

On each trial, a fixation cross appeared at the center of the screen for 1000 ms, after which two different sound sequences were presented separately to each ear. These two sequences were presented simultaneously, comprised the same number of pure tones, but differed in pitch. The sound sequence consisted of four sounds, separated by inter-sound intervals. Each sound was either a pure tone or a white noise burst, depending on the experimental condition. The duration of the pure tones, the white noise, and the inter-tone intervals was 200 ms, yielding a sound sequence that consistently lasted for 1400 ms. For the attended side, the last one (set-size 1), two (set-size 2), three (set-size 3), or four (set-size 4) sounds were randomly sampled without replacement from the predefined set of 23 target pure tones, and participants were required to memorize the exact pitches of these pure tones. White noise bursts were used to fill up the sequence where no pure tone was presented (i.e., in set-sizes below 4; see Figure 1). For the unattended side, the last one (set-size 1), two (set-size 2), three (set-size 3), or four (set-size 4) sounds consisted of distractor sounds. The distractor sound was either the lowest (125 Hz) or the highest frequency (8.1 kHz) from the predefined distractor set of pure tones (with equal probability), and was repeated once (in set-size 1 condition), twice (in set-size 2 condition), three times (in set-size 3 condition), or four times (in set-size 4 condition), matching the number of pure tones on the attended side. Similarly to stimulus presentation in the attended side, white noise bursts were used to fill up the sequence of pure tones in the unattended side (i.e., in set-sizes below 4). After the presentation of the sound sequence, a maintenance period of 2000 ms followed, during which participants had to hold the pitches in memory (1 to 4 target sounds). Finally, after the maintenance period, two probe sounds were presented for 200 ms, separately to the attended and unattended side. The probe sound on the attended side was the test sound, which could either be present (50% trials) or absent (50% trials) in the memorized sound sequence. On target-present trials, the probe sound was randomly selected from the memorized sound sequence with equal probability. On target-absent trials, a tone from the memorized sound sequence was similarly selected, but substituted with an adjacent pure tone (i.e., one step higher or one step lower in the pre-generated target set of 23 pure tones). The sound on the unattended side was the same distractor that was presented in the original sound sequence (125 Hz or 8.1 kHz). After the offset of the probe sound, participants were required to indicate as accurately as possible whether the probe sound on the attended side was present or absent in the memorized sound sequence, while ignoring the sound on the unattended side. After the response (present vs. absent), the next trial started (See Figure 1). Overall, participants completed a total of 416 trials divided into eight blocks; two practice blocks of 16 trials each, followed by six experimental blocks of 64 trials each.

**Figure 1.**
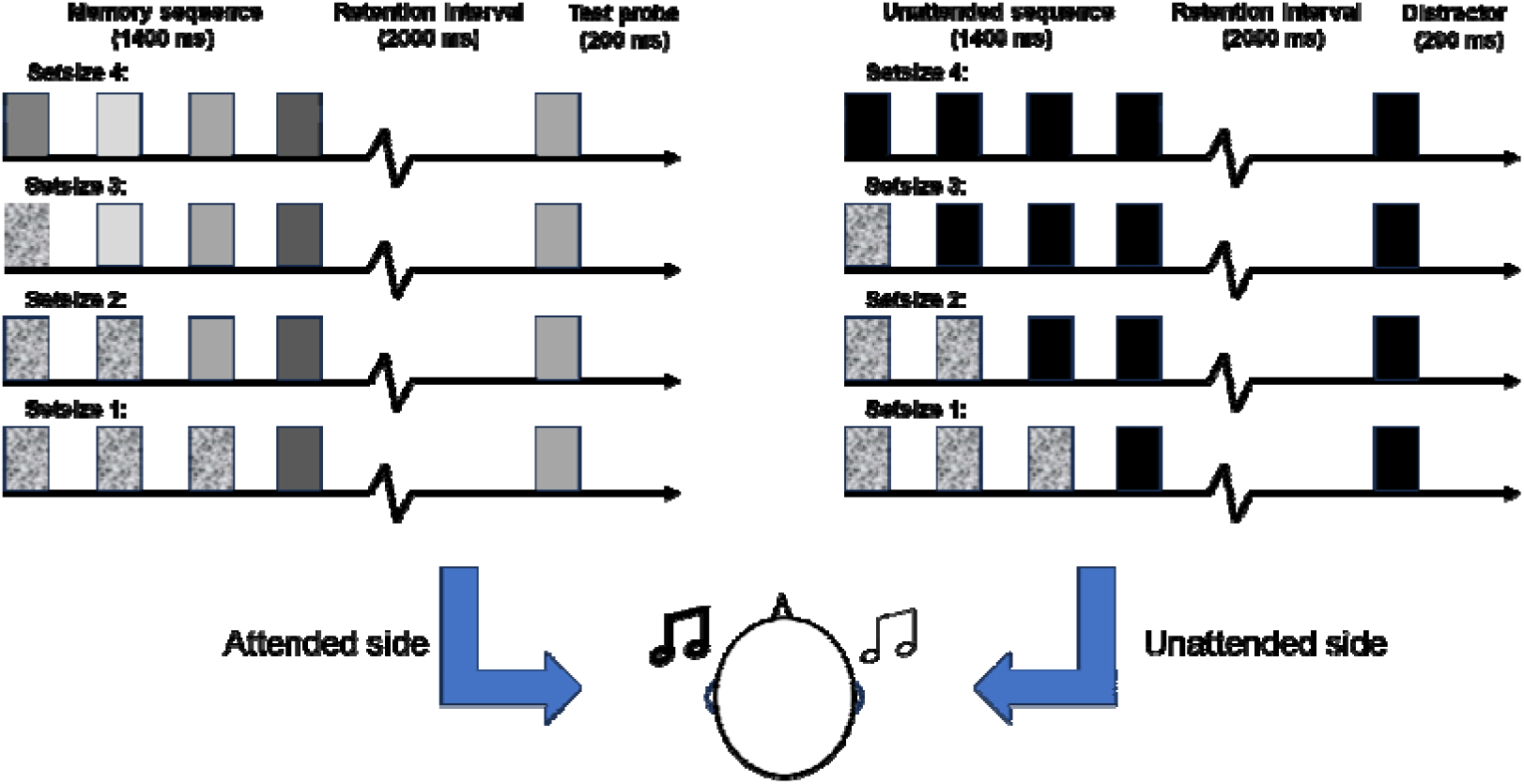
Schematic depiction of the task. Two different sound sequences were presented separately and simultaneously to each ear, whereby the participant was instructed to attend one side and ignore the other. On the attended side, the last 1, 2, 3, or 4 sounds (depending on the set-size) of the sequence were pure tones of different frequencies that participants were instructed to memorize. On the unattended side, the last 1, 2, 3, or 4 sounds in the sequence consisted of the same (lowest or highest pitched) pure tone, which participants could ignore. On both sides, the remaining (3, 2, or 1) sounds consisted of white noise. After a two second delay, during which participants maintained 1, 2, 3 or 4 sounds in memory, a probe sound was presented to the attended side while the to-be-ignored tone was presented to the other side. Participants were required to indicate whether the probe sound was present or absent in the memorized sound sequence.

### 2.4 EEG recording

EEG was recorded at a sample rate of 2048 Hz from 32 standard electrode sites placed according to the international 10/20 system using a BioSemi EEG system. Six additional EXG flat type electrodes were used to record horizontal and vertical eye movements and provide mastoid references. During the experiment, participants were instructed to fixate on the center of the monitor and try not to make horizontal or vertical eye movements.

The offline analysis of EEG data was performed using Matlab 2022a (https://www.mathworks.com/) and eeglab 14.1.2b (Delorme & Makeig, 2004). For pre- processing, EEG data were first re-referenced to the average of all 32 channels. Then a 0.01 Hz to 40 Hz band-pass filter and a 50 Hz notch filter were applied to remove high frequency noise and electromagnetic radiation from the environment. After the filtering, EEG data were down-sampled to 256 Hz to speed up computation. Then, an infomax independent component analysis (ICA) algorithm (Bell & Sejnowski, 1995) was applied to correct the signal for eye movement artifacts. The SASICA plugin was used to automatically identify the artifact component (Chaumon et al., 2015). On average, 1 independent component was rejected per participant (*SD* = 0.71, range = 0-2)^1^. Furthermore, the EEG signals were segmented into 5500 ms epochs (-1000 ms to 4500 ms relative to the onset of the sound sequence) separately for each attended side and set-size condition. The interval of 0-200 ms prior to the onset of the sound sequence served as baseline. Finally, if the voltage value at any time point in the epoch on any of the 32 channels exceeds ±200 µV, this epoch was excluded from further analysis. Six participants were excluded because an excessive number (>25%) of EEG epochs were removed (i.e., voltages exceeding ±200 μV). The average epoch rejection rate of the remaining participants was 6.40% of all trials (*SD* = 6.67%, range = 0-24.7%)^2^.

### 2.5 Behavioral data analyses

Accuracy rate and working memory capacity *K* were calculated separately for each participant in each experimental condition (Equation 1; Rouder et al., 2011). In the equation, *K* is WM capacity, *N* is the number of items in the memory sequence (set-size), and *H* and *F* are the observed hit rates and false alarms in Set-size N condition, respectively. Moreover, participants whose accuracy and or WM capacity K exceeded ±3 *SD* of the group mean were identified as outliers and removed. The ±3 *SD* threshold is commonly used in EEG and behavioral research to identify outliers that may reflect misunderstanding, disengagement, or atypical strategies (Berger & Kiefer, 2021; Mohanathasan et al., 2024). In the present study, this criterion led to exclusion of data from one participant.

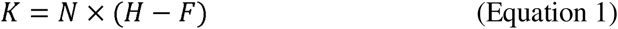

Bayesian repeated measures analyses of variance (RM ANOVAs) were conducted separately for mean accuracy and mean auditory WM capacity *K* with factors Set-size (1, 2, 3, vs. 4) and Attended side (Left, vs. Right) using JASP 0.17.3.0 (JASP Team, 2024). The default priors were applied while the seed value was consistently set to 1 for repeatability.

Moreover, effects were evaluated across matched models (Mathôt, 2017), by comparing models containing the factor of interest to equivalent models without this factor. This approach provides an “Inclusion Bayes Factor” (BF_incl_), which reflects the amount of evidence for or against the specific (main or interaction) effect of interest. We followed the guidelines suggested by Kass and Raftery (1995) for the interpretation of Bayes factors. Specifically, BFs larger than 3 signified substantial (or more) evidence in favor of the effect; BFs between 0.3 and 3 signified no conclusive evidence in favor of or against the effect; BFs smaller than 0.3 signified substantial (or more) evidence against an effect.

### 2.6 EEG Data analyses

For EEG data analyses, we aimed to answer the following two questions: (1) Is auditory WM load reflected in lateralized neural responses, akin to the CDA for visual WM load? (2) Can we identify non-lateralized load-dependent neural markers of auditory WM? To answer question 1, we computed the lateralized (i.e., contralateral minus ipsilateral) EEG response amplitude and the lateralized alpha-band (8-12 Hz) power evoked in the four different set-size conditions. For question 2, we applied multivariate decoding to patterns of alpha-band power across the scalp, as measured during the maintenance period. The focus on alpha-band over other frequency bands followed from previous studies showing close relationship between alpha oscillations and WM maintenance (Leiberg et al., 2006; Wilsch & Obleser, 2016). All data analyses were performed in Matlab with Fieldtrip toolbox (Oostenveld et al., 2011; Donders Institute for Brain, Cognition and Behaviour, Radboud University, the Netherlands. See http://fieldtriptoolbox.org) and MVPA-light toolbox (Treder, 2020).

#### 2.6.1 Lateralized responses in time and frequency domain

To test whether there is an overall lateralized (CDA-like) response (reflecting the instruction to either memorize tones on the left or the right side), contralateral and ipsilateral EEG responses were averaged across all set-size conditions. Then a cluster-based permutation test (paired *t*) (Maris & Oostenveld, 2007) was conducted to test for a difference between the contralateral and ipsilateral EEG responses during the maintenance period (1400 to 3400 ms relative to the onset of the sound sequence). This permutation test was performed on all lateralized electrodes, to assess which timepoints exhibit a CDA-like response to auditory WM load. The permutation test consisted of four steps: (1) Paired-sample *t*-tests were used to test at each time point and each electrode whether the contralateral and ipsilateral responses differed at the group level. (2) Clusters were defined as contiguous time points for which the *t*-test was significant (*p* < .05) at least at one electrode. Then, for each of these clusters, the *t*- values were summed to obtain a cluster-level *t* mass. (3) We constructed a null distribution of cluster-level *t* mass values, by randomly swapping the condition label (contralateral or ipsilateral) across trials, for each participant, and then calculating the maximum cluster-level *t* mass at the group level. Importantly, labels were swapped for an entire trial (i.e., time-series) rather than timepoint-by-timepoint, to preserve autocorrelations between timepoints in the null data. By repeating this procedure 1000 times, we obtained a null distribution of maximum cluster-level *t* mass values. (4) Finally, we computed *p* values for each of the clusters in the observed data, by computing the fraction of permuted data sets containing *t* mass values at least as extreme as that of each observed cluster (using a two-tailed alpha = 0.05). For each significant cluster, we also reported the averaged Cohen’s *d* over a rectangular region covering the cluster as effect size (Meyer et al., 2021). We considered channels to exhibit a CDA-like response, whenever channels yielded a significant lateralized response throughout the entire cluster.

To test whether the CDA-like response scaled with set-size, we followed the same procedure as above, but for each set-size condition individually. Another cluster-based permutation test was conducted to compare the lateralized responses across four set-size conditions during the maintenance period (1400 to 3400 ms relative to the onset of the sound sequence). This permutation test was performed on all lateralized electrodes, to assess during which timepoints and for which electrodes the CDA-like response scales with set-size.

A similar approach was followed to test for lateralized load-dependent responses in the time-frequency domain (alpha power). Instead of using the raw (i.e., voltage) EEG responses, a time-frequency analysis was performed first, using 7-cycle Morlet wavelet decomposition (i.e., mf0σt=7; Roach & Mathalon, 2008) for frequencies ranging between 4 and 30 Hz in 1 Hz steps. The Morlet filtering was performed by convolving single-trial EEG epochs from each scalp electrode with complex Morlet wavelets. A 7-cycle Morlet wavelet was used in this analysis to provide a good tradeoff between time and frequency resolution. The decomposition was performed on 5.5 s EEG epochs after the pre-processing described above, ranging from -1000 ms to 4500 ms relative to the onset of the sound sequence. After Morlet wavelet decomposition of the trials was performed, oscillatory power was calculated, by taking the square of the modulus of the resulting complex number. Each trial was baseline corrected using a decibel (dB) normalization, with the -200-0 ms window before onset of the sound sequence serving as baseline. Following the approach of previous studies, we focused on the power in the alpha frequency band (8-12 Hz) (Leiberg et al., 2006; Wilsch & Obleser, 2016).

Cluster-based permutation testing (following the same procedures as above) was performed to test for the existence of a lateralized alpha response, and to test whether this lateralized alpha response scales with set-size.

#### 2.6.2 EEG decoding analyses

The goal of the decoding analyses was to test for the existence of non-lateralized EEG responses that scale with auditory WM load. Compared to univariate approaches, multivariate decoding is more sensitive to uncover differences between conditions as it leverages the scalp distribution of neural signals, allowing for the detection of discriminable patterns that may not be evident in individual electrodes (Peelen & Downing, 2023). Here, we set out to classify different load conditions based on the scalp distribution of alpha-band power, using the MVPA-light toolbox (Treder, 2020) in Matlab.

We performed two distinct MVPAs to test whether we could distinguish the patterns of activity evoked by different set-size conditions. First, we trained classifiers to differentiate between set-size conditions by performing timepoint-by-timepoint decoding (i.e., training and testing the classifier on data from the same timepoint within a trial). Six different decoding comparisons (Set-size 1 vs 2, 1 vs 3, 1 vs 4, 2 vs 3, 2 vs 4, 3 vs 4) were performed separately for each time point from the onset of the sound sequence to the end of the maintenance period. Second, we conducted a temporal generalization analysis, whereby classifiers were trained and tested on all possible combinations of timepoints. This temporal generalization analysis allowed us to identify whether the neural response to auditory WM load was stable over time or was dynamically changing during the maintenance period. Here as well, we conducted six different decoding comparisons (Set-size 1 vs 2, 1 vs 3, 1 vs 4, 2 vs 3, 2 vs 4, 3 vs 4). Finally, for both analyses, we also calculated a searchlight analyses. The significant decoding accuracy for each electrode in the searchlight analyses would inform us of which electrodes are most informative to distinguish between set-size conditions.

#### Decoding: timepoint-by-timepoint classification

To test whether scalp patterns of alpha oscillations tracked auditory WM load, we conducted a timepoint-by-timepoint decoding analysis, ranging from the start of stimulus encoding to the end of the maintenance period, using electrodes as features. To obtain the mean alpha-band power, the data was first down-sampled over time by averaging every five consecutive time points (20 ms) without overlap. Then the data was z-scored, and data from each set of 5 consecutive trials within the same set-size condition were averaged to improve the signal-to-noise ratio. This resulted in 35 compound ‘trials’ (samples) on average (*SD* = 2.62, range = 28-38) that could be used for our decoding analyses.

We used linear SVM classifiers with a leave-one-trial-out cross-validation procedure to decode between set-sizes. Taking set-size 1 vs. 2 as an example, we performed classification analyses to investigate whether the scalp pattern of alpha-band power could distinguish between set-size 1 and 2 conditions. Above-chance classification would imply that the patterns of alpha-band power were load-dependent. Specifically, for a given timepoint, and on each cross-validation iteration, one compound trial (i.e., the average of 5 trials) was drawn from the set-size 1 or 2 condition. The remaining compound trials were then used to train a linear SVM classifier to distinguish between activity evoked in set-size 1 and set-size 2 conditions. This classifier was then tested on the trial that was left out of the training procedure. To keep the number of training examples equal between the two conditions, a random selection of trials was removed from the condition with the most trials. The classification was performed repeatedly until each compound trial was used to test the classifier once, thus yielding one classifier outcome per compound trial. These classifier outcomes were averaged to yield a single decoding accuracy score for a given participant and a given timepoint. This procedure was repeated for every timepoint of a trial, and for every participant, and for classifying between each of six possible pairs of set-size condition. In a final step, decoding accuracies from all six classification pairs were averaged (per timepoint) to obtain an overall classification performance for set-size.

#### Decoding: temporal generalization

To test whether the specific patterns of alpha-band power reflecting the different set-size conditions were stable or dynamic (i.e., varying over time), we performed a temporal generalization decoding approach. The procedure was identical to that of the timepoint-by-timepoint decoding describe above, with the exception that the linear SVM trained at one time point was not only tested on the same time point at which it was trained, but also tested on all other time points (King & Dehaene, 2014). Thus, the temporal generalization analyses resulted in a 2-dimensional matrix of decoding accuracies, wherein each timepoint in a trial is used for training and for testing the classifier. If the pattern of alpha-band power reflecting specific set-sizes is dynamically changing over time, significant decoding should be found only along the diagonal line (which is the same data as the timepoint-by-timepoint classification analysis described above). If the alpha oscillation pattern underlying classification performance is stable throughout the delay interval, significant decoding should not only be found along the diagonal line, but also spreading over all combinations of training and testing times during the maintenance period, resulting in a square-like shape of significant decoding performance.

#### Decoding: statistics

To test for significance in the timepoint-by-timepoint decoding analysis, we followed the same cluster-based permutation approach as described for the lateralized univariate responses above. In this case, cluster-based permutation tests were applied to compare decoding accuracy against chance level (0.5 for binary classification). The cluster-level *t* mass was computed across one dimension (i.e., timepoints within a trial). This permutation test was performed for each of the six set-size comparisons as well as the average of all set-size conditions. For the temporal generalization analysis, the same cluster-based permutation approach was used, except that the cluster-level *t* mass was now computed across two dimensions (i.e., train and test timepoints within a trial).

#### Decoding: searchlight analyses

To identify which electrodes were most informative for distinguishing between set-size conditions, we conducted an independent searchlight analysis across all significant decoding time points during the maintenance period. The rationale was that electrodes can only be considered informative when decoding accuracy is significantly above chance. Specifically, for the set-size 1 vs. 2, 1 vs. 3, and 1 vs. 4 comparisons, we defined for each electrode a cluster consisting of its neighboring electrodes. Instead of using the entire scalp pattern, we used these local clusters as features and performed timepoint-by-timepoint decoding as well as temporal generalization analyses for each participant and each significant time point. Decoding accuracies were then averaged across the significant set-size comparisons and across the significant decoding time points within the maintenance period. Finally, group-level topographies of decoding accuracy were obtained by averaging electrode-wise accuracies across participants. This procedure yielded two topographic maps showing which electrodes better distinguished between different set-size conditions during maintenance period, separately for timepoint-by-timepoint and temporal generalization decoding.

To test which electrodes were most informative for distinguishing between set-size conditions, we performed a permutation test. On each of 1000 permutations, we randomly shuffled the searchlight decoding accuracy across electrodes for each participant and each significant time point, and then applied the exact same analysis steps as described above for the observed data. For each electrode, we then computed the proportion of permutations containing any searchlight decoding accuracy at least as high as the searchlight decoding accuracy for that electrode in the observed data. These proportions were interpreted as *p*- values that implicitly account for multiple comparisons (i.e., across electrodes).

## 3 Results

### 3.1 Behavioral data

For accuracy, we found decisive evidence (*BF_incl_* = 7.85 x 10^26^) in favor of a main effect of Set-size (Figure 2a). Subsequent pairwise analyses, provided overwhelming evidence that accuracy differed between all set-size conditions (all *BF_incl_* > 2.59 x 10^5^), showing that accuracy dropped dramatically as the set-size increased. This indicates that the manipulation of auditory WM load was successful. We found no evidence for a main effect of Attended side (*BF_incl_* =.73), and substantial evidence against an interaction of Set-size and Attended side (*BF_incl_* = .14). Thus, accuracy did not systematically depend on left versus right presentation of the memory array.

**Figure 2.**
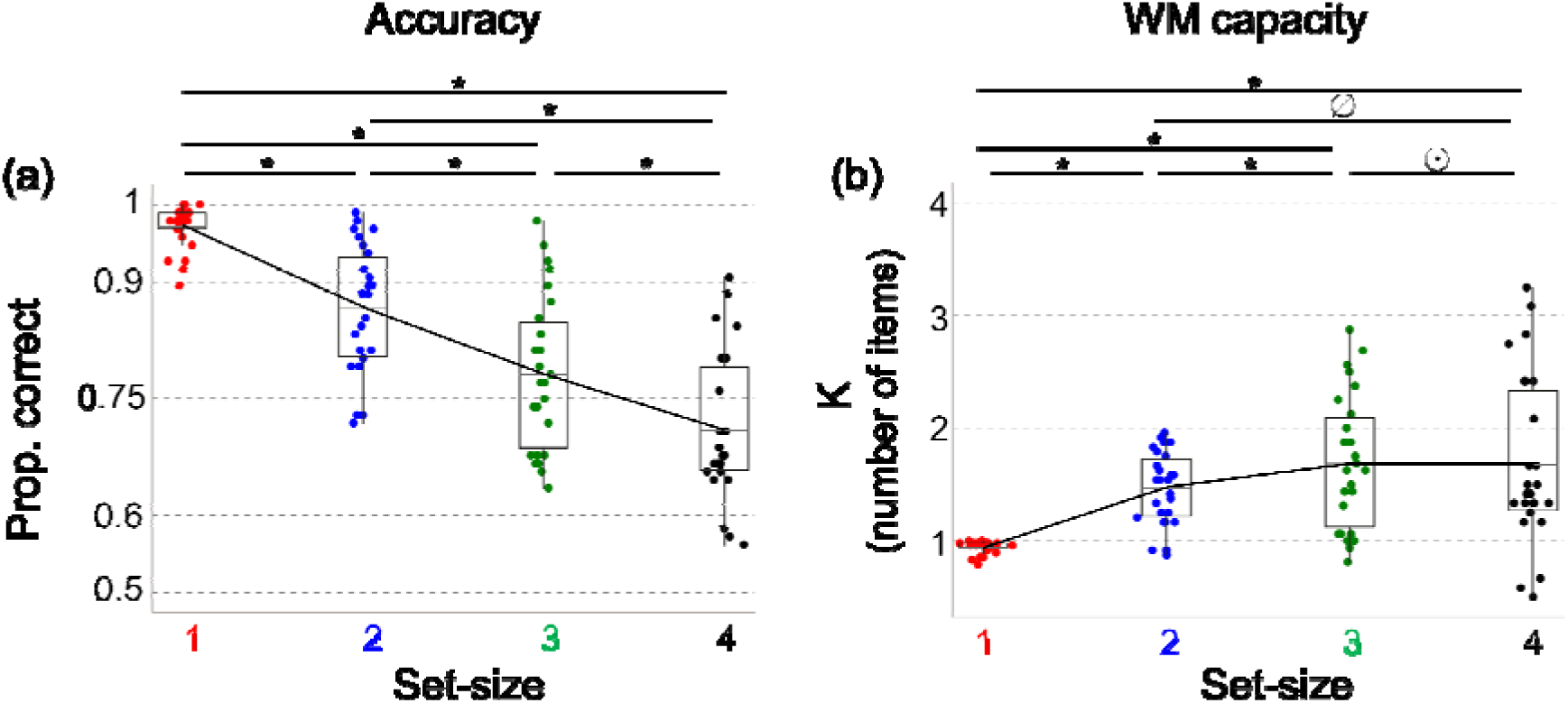
Results of behavioral performance. Panel (a) shows the accuracy, and panel (b) shows the auditory working memory capacity estimate *K,* separately for set-size 1 (red), 2 (blue), 3 (green) and 4 (black) conditions. In all plots, the mid-black-line in the box plot represents the group mean. * indicates evidence in favor of the post-hoc differences; ø indicates inconclusive evidence (1/3 < BF_10_ < 3); ⊙ indicates evidence against the post-hoc difference.

For working memory capacity, we also found decisive evidence (*BF_incl_* = 2.68 x 10^8^) in favor of a main effect of Set-size (Figure 2b). In follow-up analyses, we found evidence that WM capacity was lower in Set-size 1 than in Set-sizes 2, 3, and 4 (all *BF_incl_* > 2.52 x 10^5^), and that WM capacity was lower in Set-size 2 than in Set-size 3 (BF = 76.87). We found no conclusive evidence that WM capacity differed between Set-sizes 2 and 4 (*BF_incl_* = 1.47), and we found evidence against a difference in WM capacity between Set-sizes 3 and 4 (*BF_incl_* = .15). Finally, we found no conclusive evidence for a main effect of Attended side (*BF_incl_* = .41), and we found evidence against an interaction of Set-size and Attended side (= .09).

Together, these behavioral results demonstrate that estimated WM capacity initially increased as the set-size increased, but then leveled off when set-size (the number of items to be memorized) reached 3, and the corresponding WM capacity *K* equaled about 2 items. The relatively low capacity for auditory WM is largely consistent with previous studies (Alunni- Menichinin et al., 2014; Li et al., 2013; Prosser, 1995), in which researchers also found the maximum capacity of auditory WM was 2.8, 2.9 and 2 pure tones, respectively. Importantly, this plateau establishes that the ceiling of auditory WM capacity (in the present experimental setting) is somewhere between two to three pitches of pure tones, and indicates that our higher set-size condition(s) exceeded capacity limitations.

### 3.2 Lateralized univariate responses

To test whether auditory working memory load was reflected in lateralized neural responses, akin to the CDA for visual WM load, we calculated interhemispheric difference for each set- size condition for both mean univariate responses (time domain) and mean alpha-band power (frequency domain) during the maintenance interval. Two cluster-based permutation tests were performed to test (1) whether an overall lateralized response is observed during the maintenance period, reflecting the side of the attended memory items, and (2) whether this putative lateralized response scales with set-size.

Testing for a lateralized response across set-sizes in the time domain revealed a significant cluster across the whole maintenance period (between 1400 and 3400 ms after stimulus onset), with stronger contralateral compared to ipsilateral activity (*t*-mass = -6.80 X 10^3^, *p* < .01). The average effect size over a rectangular region spanning 10 channel pairs and 1400–3400 ms was *d* = -.59. This negative difference waves confirms the existence of a CDA-like response during the maintenance interval, indicating the validity of our attention manipulation (Figure 3a). This lateralized response, however, did not differ between set-size conditions (no significant clusters were observed), suggesting that auditory WM load is not reflected in lateralized responses.

**Figure 3.**
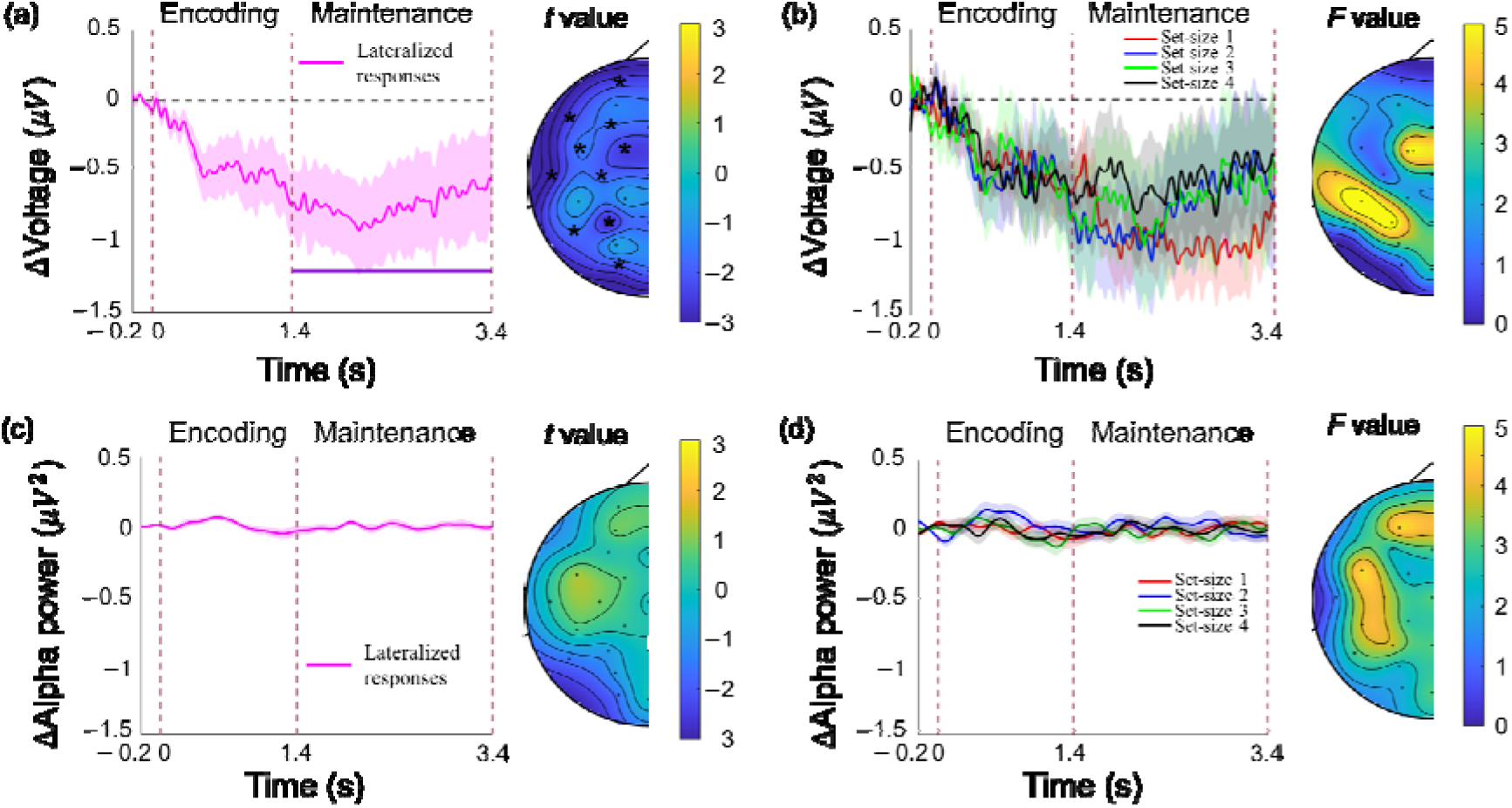
Lateralized (contralateral minus ipsilateral) response, and its topographical distribution. (a) Left: Grand average lateralized response in the time domain data averaged across all set-size conditions and all electrodes, shown from baseline (0.2 ms) until the end of retention (3.4s). The shaded areas depict the standard error of the mean. The vertical dashed lines indicate (from left to right) the onset of the tone sequence, the onset of the maintenance period, and the end of the maintenance period. The horizontal purple line indicates a significant lateralized effect (deviating from 0) collapsed across all four set-size conditions. Right: The topographical maps depict the magnitude of the overall lateralized effect (*t*-value) averaged across four set-size conditions during the time window with significant lateralized effects. Electrodes where these effects are significant are marked with an * (if any). (b) Left: Lateralized responses in the time domain data measured in the set-size 1 (red), 2 (blue), 3 (green), and 4 (black) conditions, averaged across all electrodes. Right: The topographical maps depict the magnitude of the set-size effect (*F*-value) averaged across four set-size conditions during the time window with significant lateralized effects. Panel (c) and (d) depict the same as Panel (a) and (b), but for lateralized alpha-band power.

Conducting the same two analyses with alpha-band power (in the frequency domain) revealed no significant clusters (Figure 3b); neither when collapsing across set-sizes, nor when testing for differences between set-sizes.

In sum, substantial lateralized responses were found in the time domain, reflecting which side was attended for the memory task. This demonstrates that our attention manipulation (attend left or attend right) was successful. However, we found no evidence that auditory WM load was reflected in lateralized responses, neither in the time nor in the frequency domain, suggesting that the CDA is a modality-specific rather than supra-modal marker for visual WM load only.

### 3.3 Alpha-band power decoding

#### Timepoint-by-timepoint classification

We set out to test whether WM load could be decoded from scalp patterns of EEG alpha-band oscillations. To this end, we trained a linear SVM classifier to distinguish between patterns of alpha-band power evoked by different set-size conditions. For this first decoding analysis, the classifier was trained and tested using data from the same time point in a trial (i.e., timepoint-by-timepoint decoding), ranging from the onset of the memory sequence to the end of maintenance period. The results of the cluster-based permutation test showed that, overall, set-size could be reliably decoded from alpha-band power from around 450 ms after the onset of the memory sequence, until around 3000 ms, thereby covering most of the encoding and maintenance period (1 cluster, *p* < .001, d_mean_ = .78 ; see Figure 4). The independent searchlight analyses showed that the centro-parietal electrodes were most informative to the decoding of auditory WM load during the maintenance periods. Thus, auditory WM load can be decoded from scalp patterns of alpha-band power.

**Figure 4.**
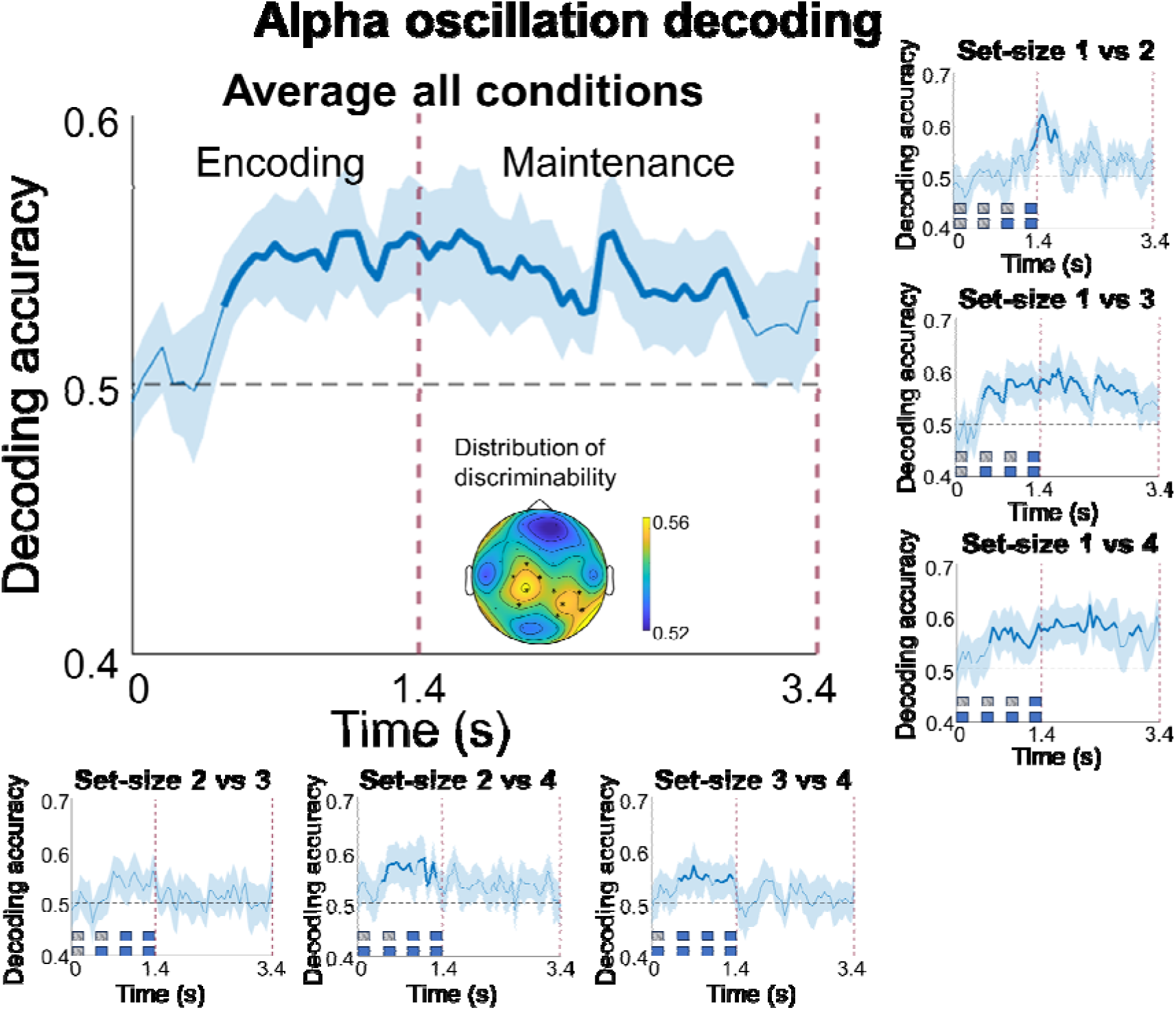
Timepoint-by-timepoint decoding of scalp patterns of alpha-band (8-12 Hz) power. All panels depict decoding accuracy (y-axis) as a function of time (x-axis). The big panel depicts decoding accuracy averaged across all six pairwise comparisons (i.e., main effect set-size), which are depicted individually in the surrounding smaller panels. Bold blue lines indicate significant above chance (50%) decoding, based on cluster- based permutation testing to account for multiple comparisons. The vertical dashed-purple lines split time into the encoding and maintenance periods. The topographical map depicts the most informative sites for distinguishing between set-size conditions during the maintenance period, based on a separate searchlight analysis. Electrodes with significant information for distinguishing between set-size conditions are marked with an *. In the small panels, the blue squares indicate the target tones, while the blurred squares indicate the white noise.

We then investigated for each pair of set-size conditions individually, whether they evoked discriminable scalp patterns of alpha-band power (i.e., 1 vs 2, 1 vs 3, 1 vs 4, 2 vs 3, 2 vs 4, and 3 vs 4). As can be seen in the six smaller panels of Figure 4, significant decoding during the maintenance period was found for the set-size 1 versus 2 (1300 ms to 1750 ms; 1 cluster, *p* < .001, *d_mean_*  = .71), set-size 1 versus 3 (1400 ms to 2200 ms, and 2300 ms to 3000 ms; 2 clusters, both *p* < .01, *d_mean_*  = .66 and .60), and set-size 1 versus 4 (1400 ms to 2650 ms, and 2850 ms to 3050 ms; 2 clusters, both *p* < .05, *d_mean_*  = .78 and .74). A visualization of alpha power patterns during the maintenance period is shown in Figure 5. These results showed that patterns of alpha-band oscillations during the maintenance period can distinguish between set-sizes of 1 and higher, but not between set-sizes of 2 and higher (which are above the group-level capacity estimations).

**Figure 5.**
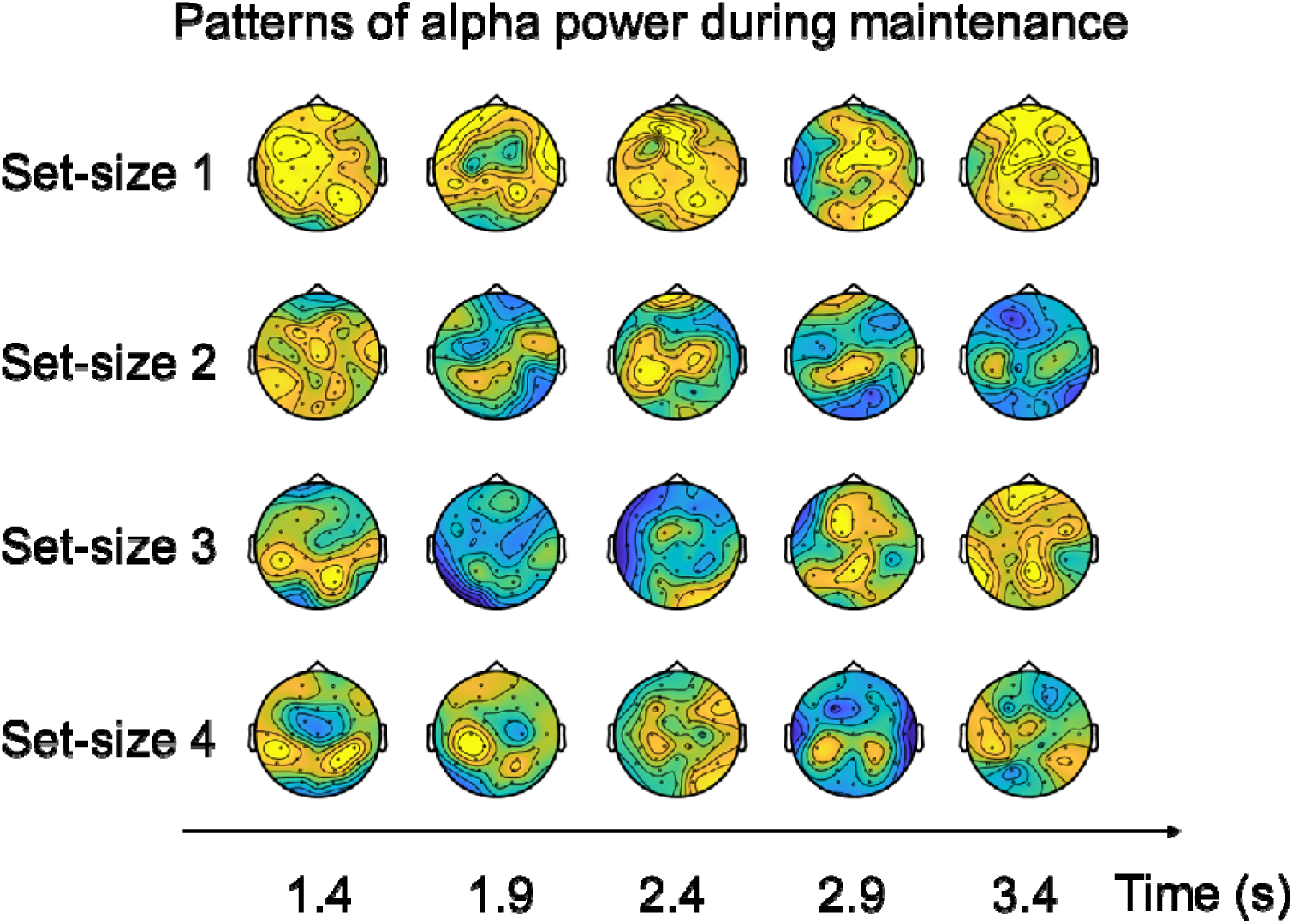
Scalp topographies of alpha-band (8–12 Hz) power during the maintenance period for set-size 1, 2, 3, and 4 conditions. Each row corresponds to one set-size condition, and each column shows a topographic map averaged over a 200-ms time window within the maintenance period.

#### Temporal generalization

In a final set of analyses, we set out to test whether the scalp patterns of alpha-band power reflecting auditory WM load are stable, or vary dynamically over the course of the maintenance period. To this end, we performed temporal generalization analyses, whereby linear SVM classifiers were trained and tested on data from all possible combinations of timepoints (thus yielding a 2D decoding matrix, instead of a time-series). When considering all set-sizes together, we replicated the finding that set-size can be decoded from scalp patterns of alpha-band power during the maintenance period (from 400 ms to 3400 ms, 1 cluster, *p* < .001,). Importantly, the present analysis also shows that significant set-size decoding (depicted in black in Figure 6) is way more prevalent on-diagonal than off-diagonal, indicating that the specific pattern of alpha-band power associated with different set-sizes varies over the course of the maintenance period.

**Figure 6.**
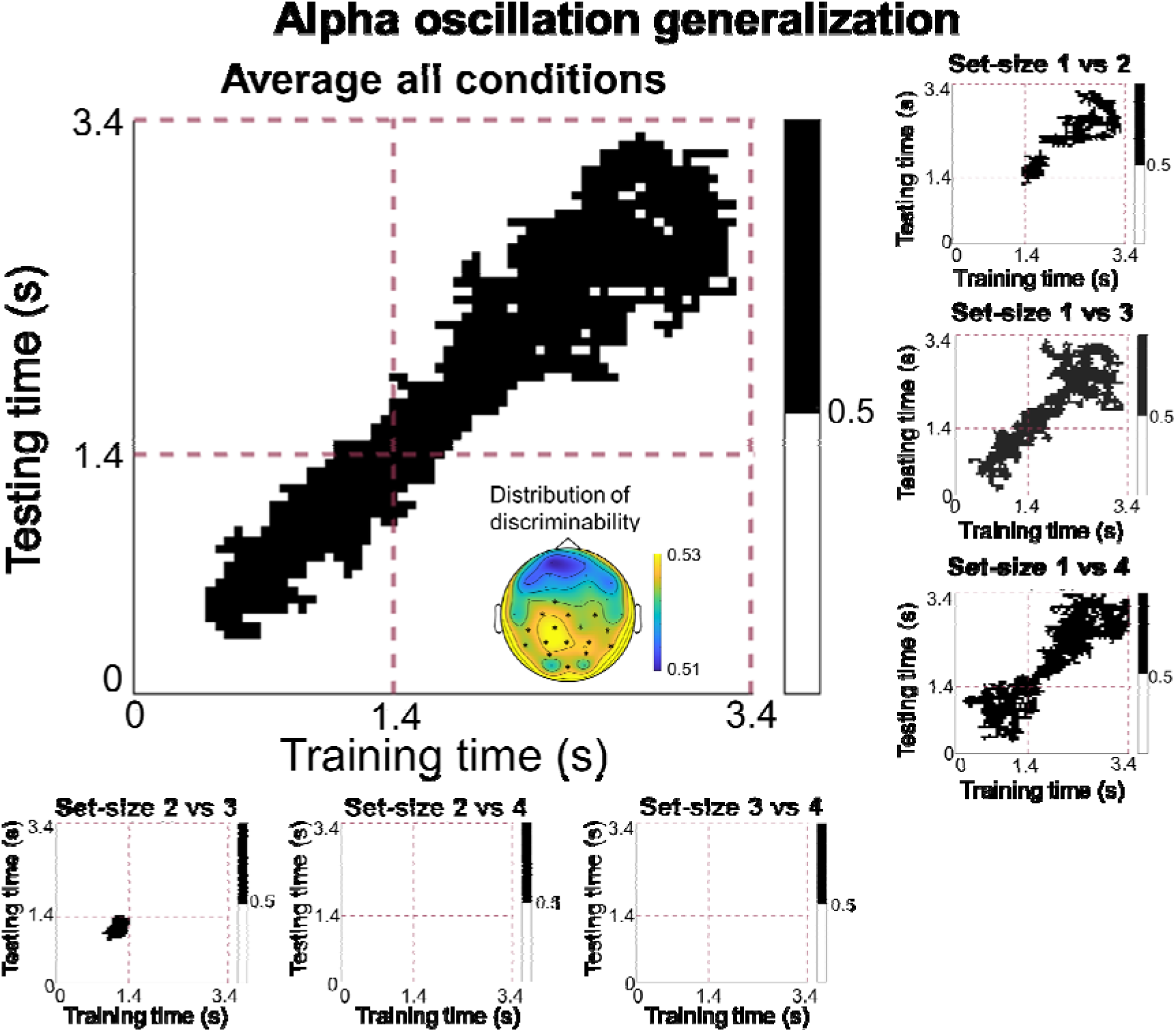
Temporal generalization decoding of scalp patterns of alpha-band (8-12 Hz) power. The big panel depicts decoding accuracy averaged across all six pairwise comparisons (i.e., main effect set-size), which are depicted individually in the surrounding smaller panels. In each plot, each data point corresponds to a classification analysis performed with training data from one time point (x-axis) and tested on another time point (y-axis). Black data points in each plot indicate significant above chance (50%) decoding, based on cluster-based permutation testing to account for multiple comparisons. The dashed-purple lines split time into encoding and maintenance periods. The topographical map depicts the most informative sites for distinguishing between set-size conditions during the maintenance period, based on a separate searchlight analysis. Electrodes with significant information for distinguishing between set-size conditions are marked with an *.

These findings are largely confirmed when considering the six set-size comparisons in isolation. As shown in the six smaller panels, temporal generalization decoding during the maintenance period revealed significant decoding for set-sizes 1 versus 2 (1400 ms to 1950 ms; 1700 ms to 3200 ms, 2 clusters, both *p* < .05, *d_mean_*  = .46 and .36), 1 versus 3 (1400 ms to 3250 ms, 1 cluster, *p* < .001, *d_mean_*  = .38), and set-size 1 versus 4 (1450 ms to 3350 ms, 1 cluster, *p* < .001, *d_mean_*  = .38). Again, no significant decoding performance was found for set-sizes 2 versus 3, 2 versus 4, and 3 versus 4, thus mirroring the results of the timepoint-by-timepoint decoding analysis.

In sum, we found that scalp patterns of alpha-band power during the maintenance period reflects auditory WM load, with centro-parietal electrodes being most informative to distinguish between set-sizes. These load-specific EEG responses allowed to distinguish between load conditions up to the group-level capacity limitations, but not between load conditions exceeding this limit, as established from the behavioral data (i.e., around K = 2). Interestingly, the specific patterns of alpha-band power that reflected specific set-size conditions evolved dynamically across the maintenance period, suggesting the dynamic coding of auditory WM load.

## 4 General Discussion

In the present study, we set out to investigate whether auditory WM load is reflected in lateralized neural responses, akin to the CDA for visual WM load. We further attempted to identify other (non-lateralized) load-dependent neural markers of auditory WM, using multivariate pattern analysis (MVPA). To these aims, we recorded EEG while participants were holding the pitches of 1, 2, 3, or 4 tones in WM for a subsequent auditory recognition task. The behavioral results showed that auditory WM capacity plateaued between two and three tones. Two key findings emerged from the analyses of EEG data. First, although we identified a lateralized (CDA-like) response during the maintenance period, this response did not scale with set-size; neither in the time nor in the frequency (alpha oscillation) domain. Thus, we found no evidence that WM load was reflected in lateralized responses. Second, using MVPA, we found that scalp patterns of alpha-band power during the maintenance period reflected auditory WM load. These load-specific EEG responses were mostly confined to bilateral centro-parietal channels, and allowed to distinguish between set-size conditions up until –but not above– group level capacity limitations (i.e., about 2 items, based on the behavioral data). Interestingly, we also found that the scalp patterns of alpha-band power reflecting specific auditory WM were not stable across the maintenance period. Instead, these patterns dynamically changed across the maintenance period, with patterns evoked at the start of the maintenance period barely generalizing to patterns at the end of the interval, and vice versa. In short, our results show that dynamic scalp patterns of alpha-band power can be used as a novel neural marker of auditory WM load. In the following sections, we will first discuss what it means that (unlike for visual WM load) auditory WM load is not reflected in lateralized responses. Then, we will elaborate on the scalp patterns of alpha power that were shown to track auditory WM load, and discuss what they tell us about the neural coding of auditory WM.

### 4.1 No lateralized responses to auditory

WM load In the current study, we identified a lateralized univariate response during the WM delay, akin to the CDA for visual WM. Control analyses indicated that the observed lateralized response is unlikely to be driven by eye movements (see supplementary materials 1.1 for more details). Unlike the CDA, however, we found no evidence that the lateralized response to auditory WM was modulated by load, neither in the time nor in the frequency domain. That is, where the amplitude of the CDA increases when visual WM load increases, the lateralized responses to auditory WM load observed in our study did not differ between load conditions. This raises the question why auditory WM load is not reflected in lateralized responses, while visual WM load is, even when both require the maintenance of lateralized sensory input? We approach this question from a cognitive, an anatomical, and a functional perspective.

From a cognitive perspective, our results can be framed in the context of the multi-component model of WM, proposed by Baddeley and Hitch (1974). The absence of a lateralized response to auditory WM load suggests that the CDA as observed for visual WM may reflect load in the visuo-spatial sketchpad specifically, rather than load in the domain-general central executive or episodic buffer. Thus, although regulating the quantity of information in WM (i.e., load) may be regarded as an executive process, sustained neural responses tracking visual WM load (i.e., the CDA) may instead be more sensory-like, and depend on the specific content that is maintained in WM. This may explain why neural markers of WM load are not so much observed in frontal electrodes – that are typically associated with executive processes – but tend to arise in sensory-specific processing regions. That is, the CDA is typically most pronounced in posterior electrodes for visual WM, and we found neural responses tracking auditory WM load (scalp patterns of alpha power, discussed below) to arise predominantly around temporal electrodes. Accordingly, different modalities may rely on qualitatively distinct storage mechanisms. However, as our data are restricted to the auditory modality, further cross-modal investigations are needed to directly assess this distinction.

From an anatomical perspective, the differences between visual and auditory WM may be due to the difference in the visual and auditory processing pathways, leading from sensory receptors to the cortex. For the visual pathway, due to the optic chiasm, items presented in the left hemifield are projected to the right primary visual cortex, whereas items presented to the right hemifield are projected to the left primary visual cortex (De Moraes, 2013; Rodieck, 1979). Thus, these hemisphere specific responses during stimulus encoding may foster a difference between contralateral and ipsilateral responses during the WM delay, when items on one side are to be remembered while items on the other side are to be ignored. In the case of auditory processing, however, the input to each ear is first extensively processed and combined subcortically, before being projected to the left and right primary auditory cortex (Pickles, 2015). Specifically, the superior olivary complex receives information from both contralateral and ipsilateral ears, to support sound localization. Then the auditory information is projected to both contralateral and ipsilateral sides of the superior colliculus (Hackney, 1987), before projecting to the primary auditory cortices. Thus, for the auditory pathway, tones presented via the left or right ear activate both the left and right auditory cortex, albeit with a bias toward the contralateral auditory cortex (Lipschutz et al., 2002; Woldorff et al., 1999). This slight contralateral dominance in auditory processing might explain the overall lateralized responses that we observed during the WM delay. Importantly, however, contralateral dominance is much weaker in auditory processing than in visual processing, because auditory input from both ears is largely integrated sub-cortically (Schwartz, 1992). This reduced lateralization during auditory processing may explain the absence of a load-dependent modulation in lateralized responses for auditory WM load.

From a functional perspective, the reason for why we did not find a lateralized response to auditory WM load might stem from the different organizational principles of auditory compared to visual information. In visual WM tasks, visual information is processed via spatial coding, while in auditory WM tasks, auditory information may be processed predominantly through temporal coding. For instance, while it is known that observers use spatial location as an organizational principle to help maintain visual information (even if the spatial location is irrelevant for the task at hand (Arora et al., 2024; van Ede et al., 2019)), several studies have shown that auditory spatial information only has an effect on working memory and perception when space is relevant for the task at hand (Klatt et al., 2018a; Klatt et al., 2018b). In the current study, the spatial location of the to-be-memorized array was irrelevant for the auditory WM recognition task. In contrast to the spatial organization observed during the maintenance of visual WM items, the maintenance of auditory WM items (such as pitches of pure tones) may be organized within a temporal reference frame, by iterating through the different items over time. This interpretation fits well with the classical example of the phonological loop (the auditory counterpart of the visuo-spatial sketchpad) as the repetition of sequences of items over time, such as the digits of a to-be-memorized phone number. In this case, at any given timepoint (in a singular brain area), although load may influence the overall system state, the number of items concurrently represented in working memory at each point in time is always 1. If auditory WM items are indeed segregated over time rather than over space, then their location is unlikely to be encoded or retained, and thus a lateralized neural response to load would not be expected to scale with set-size. The finding that our neural markers of auditory WM load (scalp patterns of alpha oscillations) appear to change over the course of the maintenance period (see below), is very much in line with this idea of temporal coding. Following the same line of reasoning, we also consider it unlikely that presenting sounds from different horizontal locations on the same side would produce lateralized ERP responses that scale with set-size.

Another possible explanation is that participants may have perceptually grouped the sequential tones into a melodic structure, thereby reducing item-specific set-size effect. However, our task design discouraged such strategies by using single-tone probes, and previous studies observed load-dependent ERP effects under similar sequential presentations. Moreover, behavioral results revealed a capacity plateau at around two items, consistent with item-based working memory. Thus, while we cannot fully rule out melodic encoding, it is unlikely to be the sole factor underlying the null ERP load effect.

Notably, given the similarity between our paradigm and the original SAN studies (Alunni-Menichini et al., 2014; Lefebvre et al., 2013), we examined whether auditory working memory load was reflected in non-lateralized neural responses during the maintenance period. To this end, we tested for differences between set-size using both univariate ERP responses and multivariate decoding of scalp voltage patterns (see Supplementary Materials 1.3 for more details). In short, during the encoding period, we observe clearly distinct EEG responses to target pure tones compared to white noise distractors, which was also reflected in the distinct time windows of significant decoding across set-size comparisons (e.g., 1 vs 4, compared to 3 vs 4). Although a sustained anterior negative wave was observed during the maintenance period, both univariate and multivariate analyses revealed that EEG responses during the delay were not reliably modulated by set-size. Thus, non-lateralized response did not scale with set-size in the current study.

Taken together, the absence of both lateralized and non-lateralized ERP set-size effects during the maintenance period suggest that, at least in the present paradigm, ERP activity did not reflect abstract non-spatial working memory representations. This interpretation is consistent with the view that previously reported lateralized effects (such as CDA) may be tied to spatially specific, sensory-like storage mechanisms (e.g. Klaver et al., 1999, Talsma et al., 2001).

### 4.2 Alpha patterns reflecting auditory WM load

Using both timepoint-by-timepoint decoding and temporal generalization decoding, we found that scalp patterns of alpha-band power during the delay period allowed to distinguish between individual load conditions up until WM capacity limitations (Set-size 1 vs 2, 1 vs 3, 1 vs 4). In contrast, patterns of alpha-band power during maintenance could not distinguish between load conditions that exceeded capacity limitations based on the behavioral results (Set-size 2 vs 3, 2 vs 4, 3 vs 4). The correspondence between these behavioral results and decoding results further substantiates that patterns of alpha-band power specifically reflect auditory WM load. The differences in alpha-band power patterns between load conditions might reflect content-invariant differences in executive demands: for instance, competition between items or switching from one item to another during the delay, which scales with the number of tones held in working memory (Leiberg et al., 2006). The more tones held in memory, the more resources had to be allocated over areas where representations of the to-be-memorized stimuli are presumably stored. Importantly, the topographical maps reveal that alpha power patterns were most pronounced in the set-size 1 condition, but their overall amplitude decreased with increasing WM load (set-size 2 - 4), indicating an overall attenuation of alpha activity as more items were held in working memory. This decrease is consistent with recent accounts that interpret attenuated alpha activity as reflecting stronger recruitment of task-relevant sensory areas for mnemonic retention and attentional prioritization (Fukuda et al., 2015; van Ede, 2018). Notably, previous studies have also found alpha power increase with working memory load. In these studies, the role of alpha oscillations is interpreted to reflect top-down inhibition of task- irrelevant sensory input and/or task-irrelevant neural processes (Kaiser et al., 2007; Van Dijk et al., 2010; Wilsch & Obleser, 2016). Most of the studies, however, compared responses in a memory task to that of a non-memory control task, and found stronger alpha-band power in the memory task. Therefore, during the maintenance period in the memory task, participants had to inhibit irrelevant information for successful retention while participants did not need to do so in the non-memory control task. In the current study, we instead compared different load conditions. In these different load conditions, the number of to-be-remembered items (presented to the attended side) varied, but the number of to-be-ignored distractor items (presented to the unattended side) was constant. The set-size of the distractors, therefore, did not covary with the set-size of the memory items. As such, the differences in scalp patterns of alpha-band power between set-size conditions reflect WM load, rather than distractor inhibition. Thus, our findings reveal the role of alpha oscillations in the maintenance of information in auditory WM.

The absence of generalization between the encoding and maintenance period (in our temporal generalization analyses) indicates that our significant decoding during the maintenance period specifically reflect maintenance-related process, rather than residual signals from the encoding period. Focusing on the maintenance period, our temporal generalization analyses suggest that the scalp patterns of alpha-band power associated with specific WM loads are not stable over time, but change over the course of the maintenance period. One possible interpretation of this, is that auditory WM load is inherently represented through dynamic coding (akin to visual WM content; see Stokes, 2015). In the context of load, changing neural representations may also reflect the interplay, over time, between sensory regions involved in maintenance, and frontal regions involved in executive processes. Another, potentially related possibility, could be that participants were maintaining the different pitches sequentially (i.e., as they were presented). In this scenario, the changing patterns of alpha-band power associated with specific load conditions, may reflect the duration of the sequence or the number of consecutive items in a sequence. It should be noted, though, that in the later stages of the delay period, we did observe some off-diagonal decoding, indicating some temporal generalization of the load-specific responses. This may either reflect smearing out of the load-specific responses as they desynchronize over time, or it may reflect the stabilization of the memory content into a format that is relevant for the upcoming test (the timing of which was predictable). During most of the delay period, however, decoding of individual load conditions was substantially more pronounced on the diagonal compared to the off-diagonal. We therefore conclude that, overall, patterns of alpha-band power reflecting auditory WM load change dynamically during the maintenance period. Future research is needed to understand what cognitive processes or storage mechanisms underly these dynamics.

If neural markers of WM load (such as scalp patterns of alpha-band power) are modality specific, they would be expected to mostly involve sensory processing regions. Indeed, we found that bilateral centro-parietal electrodes were most distinguishable to the decoding of auditory WM load during the maintenance periods, rather than posterior or frontal electrodes in visual working memory studies. Previous studies have shown the involvement of temporo-parietal regions, including auditory cortex and supramarginal gyrus, in auditory WM (Gaab et al., 2003; Grimault et al., 2014; Koelsch et al., 2009). Consistently, here the searchlight analyses revealed that patterns in bilateral centro-parietal were most distinguishable across set-sizes in classification. This is in line with the sensory recruitment hypothesis (Gayet et al., 2018; Katus et al., 2015; Scimeca et al., 2018; Silvanto & Soto, 2012), which has been more extensively studied in visual WM, and proposes that the same neural populations that represent sensory features during perception are also recruited during WM maintenance, thereby reducing cortical redundancy.

Taken together, our decoding results identified alpha-band patterns as a neural marker of auditory working memory load, reflected in reduced alpha activity. Such decreases are commonly interpreted as stronger sensory recruitment, which is consistent with our searchlight analyses showing that centro-parietal channels were most informative in distinguishing set-size conditions. This interpretation is also in line with accounts of temporal coding (e.g., rehearsal) in auditory working memory, in which repeating tones are thought to be maintained within auditory cortical areas. Moreover, our temporal generalization results revealed dynamically changing load-related patterns, further supporting the view that auditory WM relies on temporally coded rehearsal of pure tones within auditory cortex, in line with the sensory recruitment hypothesis.

### 4.3 Limitations

One may argue that the capacity of auditory WM is relative low (around 2 tones) in the current study. However, this relatively low capacity is largely consistent with previous studies (Alunni-Menichinin et al., 2014; Li et al., 2013; Prosser, 1995), in which researchers also found the maximum capacity of auditory WM was 2.8, 2.9 and 2 pure tones, respectively. It should be noted that calculations of WM capacity are estimates, and do not reflect perfect measurement of WM capacity. Indeed, capacity estimates should theoretically remain constant across set-size conditions, but the estimate of WM capacity K, is known to vary when large differences in set-size are used (Rouder et al., 2011). Thus, it is unclear whether auditory WM capacity in our study should be estimated to be around 2 items (the actual capacity estimate K) or at ∼3-4 items (the set-size conditions at which the capacity estimate K started to plateau). Could the challenging dichotic presentation tasks employed in our have reduced capacity estimates? Studies using binaural stimulation typically show that observers can ignore the unattended auditory stream with little to no effort, yielding virtually no interference to the processing of the attended auditory stream (Alho et al., 1994; Carpenter et al., 2002). Thus, the use of binaural stimulation is unlikely to have substantially reduced capacity estimations in our study. Taken together, the present behavioral results cannot be taken to reflect a universal auditory WM capacity limit, but they do demonstrate the relatively low capacity as compared to (for instance) visual WM in similar task designs. ^3^

Another potential limitation is that we manipulated the to-be-attended side in a block-wise rather than trial-wise manner. The majority of visual WM studies reporting CDA effects used trial-wise cueing, which could be argued to enhance lateralized responses. We opted for a block design to minimize confusion in our challenging dichotic auditory task. Importantly, however, several studies have reported reliable CDA effects using blocked designs (e.g., Carlisle et al., 2011; Katus et al., 2017; Reinhart et al., 2014), suggesting that CDA components can be elicited and measured under such conditions. Thus, it seems unlikely that the absence of CDA effects in the current study is solely due to the blocked design.

### 4.4 Conclusion

In conclusion, we observed two main findings in the current study. First, we found no evidence that auditory WM load is reflected in lateralized responses – neither in the time nor in the frequency domain. This implies that CDA-like responses as observed for visual WM load are vision-specific rather than domain general markers of WM load. The lack of location-specific response further suggests that auditory WM is not inherently spatially organized, as is the case for visual WM. Second, using multivariate pattern analyses, we found that scalp patterns of alpha-band power during the maintenance period reflect auditory WM load. Interestingly, patterns of alpha-band power reflecting specific load conditions were changing dynamically over the course of the maintenance period, revealing that (1) principles of dynamic neural population coding –which is known to underly the storage of WM content– may also be extended to executive WM processes, and (2) auditory WM may be inherently temporally organized, reflecting the repetition of information streams over time.

## Data availability statement

All data, codes to run the experiments, analyze and visualize the data are uploaded to the OSF platform (https://osf.io/mw5ve/).

## Ethics statement

The study protocols were approved by the faculty ethics committee (FETC) of Utrecht University (number 18-048 van der Stoep). All participants signed informed consent for their participation.

## Author Contribution

Yichen Yuan: conceptualization, data curation, formal analysis, funding acquisition, methodology, resources, software, validation, visualization, writing—original draft, writing— review and editing. Surya Gayet: conceptualization, methodology, project administration, resources, supervision, writing—original draft, writing—review and editing. Derk Wisman: conceptualization, data curation, investigation, methodology, resources, software, writing— review and editing. Stefan van der Stigchel: conceptualization, methodology, project administration, resources, supervision, writing—review and editing. Nathan van der Stoep: conceptualization, methodology, project administration, resources, supervision, writing— original draft, writing—review and editing. All authors contributed to the article and approved the submitted version.

## Funding

This work was supported by the China Scholarship Council (grant number 202206380011 to Yichen Yuan).

## Footnotes

1 A more strict criterium (eye movement correction plus ADJUST) did not change the lateralized ERPs and alpha power results (see supplementary materials 1.1 for more details).

2 The grand-average ERP responses in all eight conditions can be found in supplementary material 1.2.

3 Unlike tasks where verbal labeling is possible, which can greatly increase WM capacity, participants in the present study could not rely on such a strategy. This may have contributed to the observed relatively low capacity estimates.

## Notes

https://osf.io/mw5ve/

### Competing Interest Statement

The authors have declared no competing interest.

### Summary of Updates

Main manu: an independent searchlight analyses to identify the most informative channel; replotting Figure 3; motivating our choice for a blocked design; reporting the effect sizes for cluster-based analyses; expanding our discussion of alpha modulations in both the Introduction and Discussion sections Supplementary material: non-lateralized ERPs to working memory load using both univariate and multivariate methods, the potential contribution of eye movements to the lateralized responses.

