## Supplementary material for "Decoding auditory working memory load from EEG alpha oscillations"

**Supplementary materials**

### **1.1 Eye-movements and lateralized responses**

We consider it unlikely that the overall lateralized response is driven by eye movements. Eye movement-related activity should have been identified and removed during preprocessing by means of ICAs. Figure S1 illustrates one participant’s data from a frontal channel (AF3) before and after ICA and eye-movement correction, both at the single-trial level (a) and across the entire recording (b). This shows that the ICA successfully identified and removed the eye movement components.

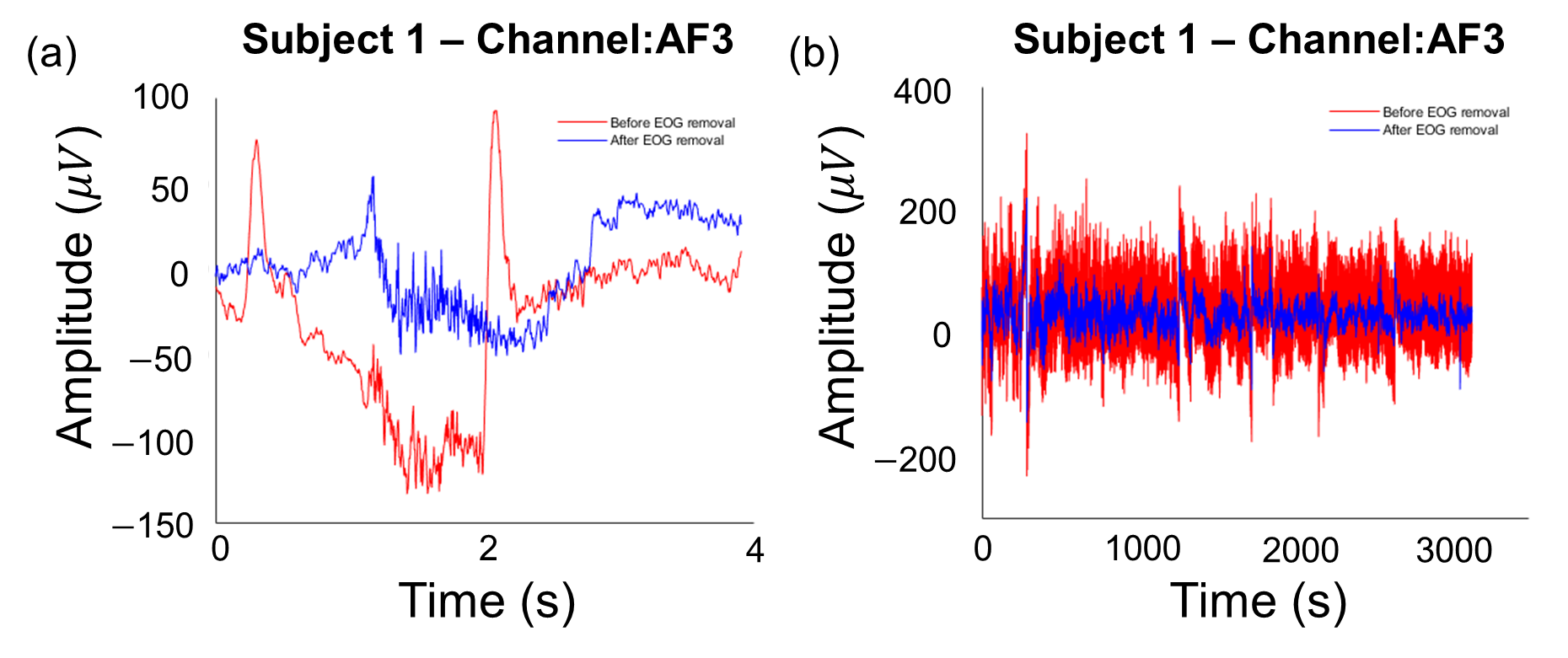

Figure S1. An example of one participants’ frontal channel data before and after ICA and eye-movement correction, both at the single-trial level (a) and across the entire recording (b).

To further exclude the possibility that the observed lateralized responses to the attended side reflected failure of the ICA to fully remove the eye movement components, we re-ran our preprocessing pipeline with only one change: using a stricter ICA and eye movement correction (eye movement correction plus ADJUST). In short, all the key results regarding lateralization remained the same: (1) we observed an overall lateralized responses in the time domain, although with shorter time window during the maintenance period compared to our original results. (2) The lateralized responses did not scale with auditory WM load, neither in the time nor in the frequency domain (Figure S2a & S2b).

Conducting the same two analyses with alpha-band power (in the frequency domain) revealed no significant clusters (Figure S2c & S2d); neither when collapsing across set-sizes, nor when testing for differences between set-sizes.

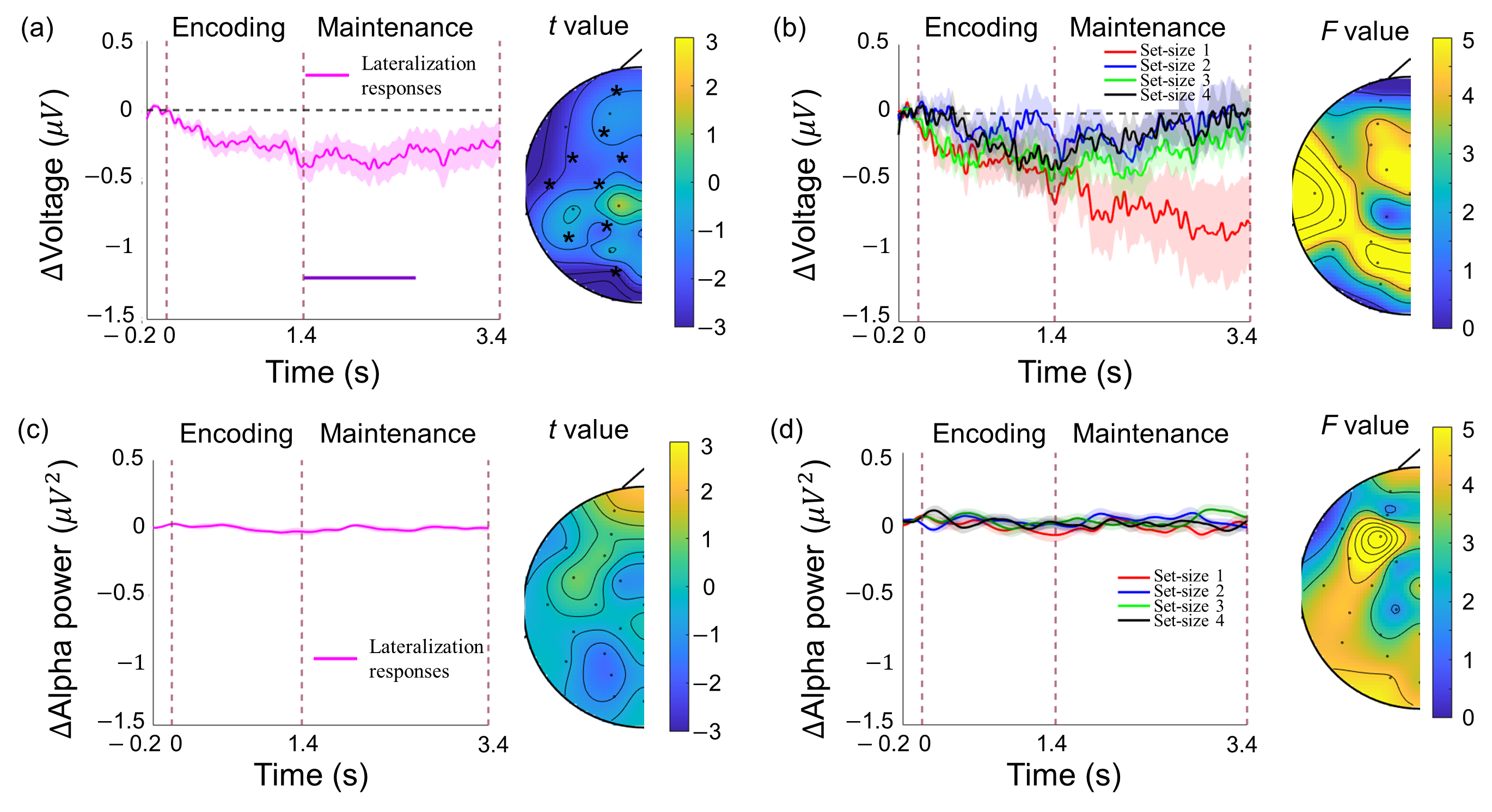

Figure S2. Lateralized (contralateral minus ipsilateral) responses, and their topographical distributions. (a) Left: Grand-average lateralized responses in the time-domain data averaged across all set-size conditions and all electrodes, shown from baseline ($-$0.2 ms) until the end of retention (3.4s). The shaded areas depict the standard error of the mean. The vertical dashed lines indicate (from left to right) the onset of the tone sequence, the onset of the maintenance period, and the end of the maintenance period. The horizontal purple line indicates a significant lateralized effect (deviating from 0) when collapsing across all four set-size conditions. Right: The topographical maps depict the magnitude of the overall lateralized effect (*t*-value) averaged across four set-size conditions during the time window with significant lateralized effects. Electrodes where these effects are significant are marked with an * (if any). (b) Left: Lateralized responses in the time-domain data measured in the Set-size 1 (red), 2 (blue), 3 (green), and 4 (black) conditions, averaged across all electrodes. Right: The topographical maps depict the magnitude of the set-size effect (*F*-value) averaged across four set-size conditions during the time window with significant lateralized effects. Panel (c) and (d) depict the same as Panel (a) and (b), but for lateralized alpha-band power.”

Regarding small saccades, although we cannot rule them out, we consider it unlikely that they are driving the lateralized responses observed here. (1) The task design provided no motivation for eye movements, as participants only viewed a fixation cross during the retention interval in this auditory WM task. (2) Even if small saccades occurred, a recent study has shown that small saccades did not reliably affect EEG responses and alpha lateralization (Arora et al., 2025).

Taken together, we consider it unlikely that the observed lateralized responses were caused (or abolished) by eye movements or small saccades.

### **1.2 Grand-average ERP responses**

To provide a comprehensive view of the raw data, we plot the grand-average ERPs across participants for all 8 conditions (2 attend-side * 4 set-size) with an extended pre-stimulus period (Figure S3). Data are shown separately for frontal (Fp1, Fp2, AF3, AF4, F7, F3, Fz, F4, F8, FC5, FC1, FC2, FC6) and posterior (CP5, CP1, CP2, CP6, P7, P3, Pz, P4, P8, PO3, PO4, O1, Oz, O2) electrode clusters.

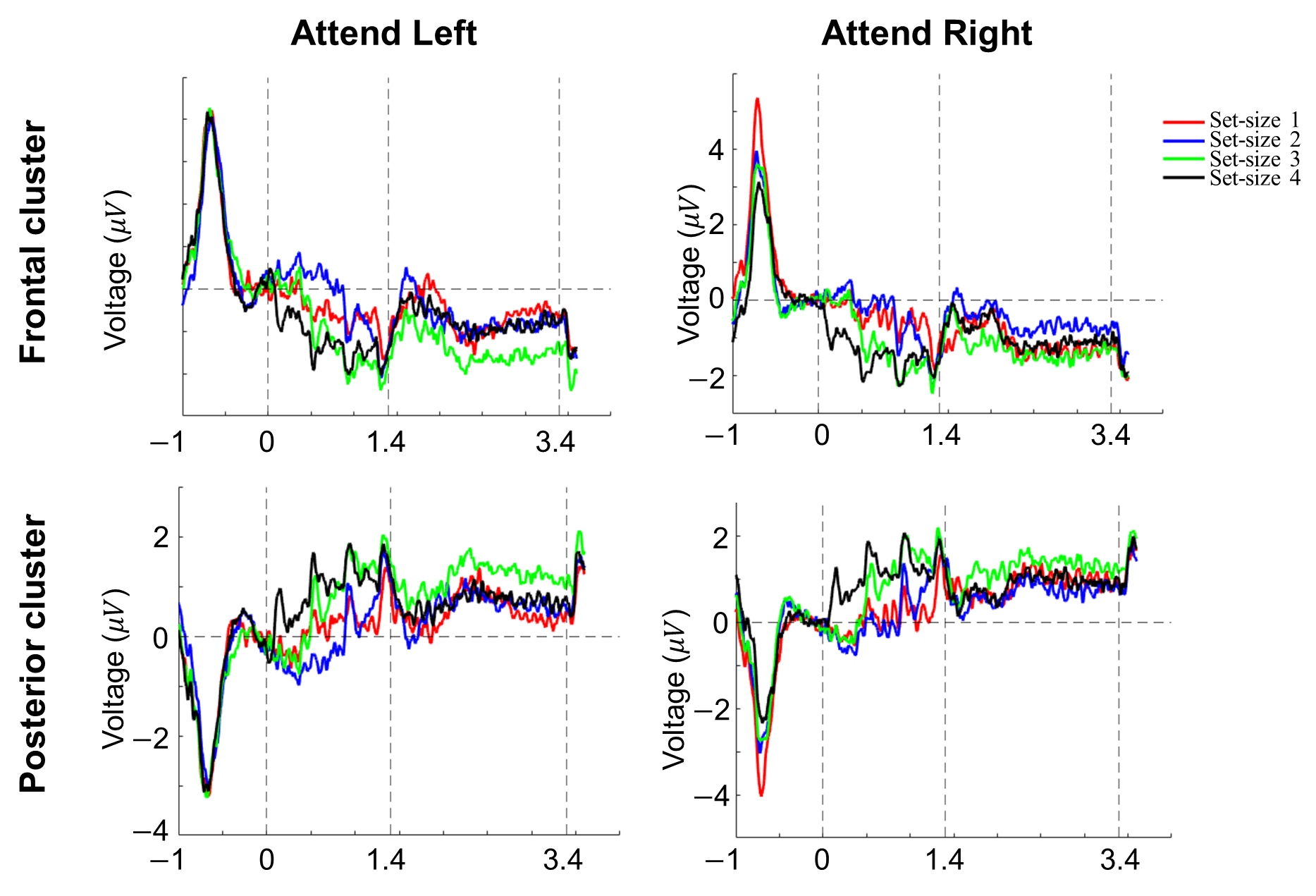

Figure S3. The grand average ERPs across participants for all 8 conditions (2 attend-side * 4 set-size) from $-$1s until the end of retention (3.6s), separated into frontal (Fp1, Fp2, AF3, AF4, F7, F3, Fz, F4, F8, FC5, FC1, FC2, FC6) and posterior (CP5, CP1, CP2, CP6, P7, P3, Pz, P4, P8, PO3, PO4, O1, Oz, O2) electrode clusters. The vertical dashed lines indicate (from left to right) the onset of the tone sequence, the onset of the maintenance period, and the end of the maintenance period.

Three aspects of the data stand out from these plots: (1) Frontal and posterior clusters showed responses in opposite polarities, likely reflecting dipole activity driven by binaural stimulation; (2) A brief but strong evoked response to the fixation cross (presented at $-$1s), which quickly returned to baseline before the onset of the sound sequence (note: this was the only visual stimulation in the trial). Importantly, a two-way repeated-measures ANOVA on the baseline period ($-$200 to 0 ms), with factors attended side (left vs. right) and set-size (1, 2, 3, vs. 4), revealed no significant main effects or interactions (all *p* > .05), confirming baseline equivalence across conditions; (3) During the encoding phase, target pure tones evoked larger responses than white noise distractors. Clear response peaks are observed in response to the pure tones relative to the noise, differentiating between set-sizes, four, three, two, and one respectively. This further substantiates the quality of the data.

### **1.3 Non-lateralized ERP responses**

Given the similarity between our paradigm and the original SAN studies (Alunni-Menichini et al., 2014; Lefebvre et al., 2013), we examined whether auditory working memory load was reflected in non-lateralized ERP responses during the maintenance period. To this end, we tested for differences between set-size using both univariate ERP responses and multivariate decoding of scalp voltage patterns.

***Univariate ERP analyses***

To test whether non-lateralized responses scale with set-size, we collapsed across attend-left and attend-right condition. Following the approach of the SAN studies, we focused on a fronto-central electrode cluster (Fp1, Fp2, AF3, AF4, F3, Fz, F4, FC1, FC2). For each participant, ERP amplitudes were averaged across these electrodes during the maintenance interval (1400–3400 ms relative to sequence onset). A repeated-measures ANOVA was then performed on mean amplitudes with factor Set-size (1, 2, 3, vs. 4). In addition, similar to the analyses in the main manuscript (section 2.6.1), we conducted cluster-based permutation tests across all electrodes to identify clusters and timepoints that scaled with set-size during the maintenance period.

Figure S4 shows non-lateralized ERPs and delay-period topographical distributions for set-size 1, 2, 3 and 4 conditions. Although a sustained anterior negative wave (resembling the SAN component) was observed in all set-size conditions, the ANOVA on mean amplitudes during the maintenance revealed no reliable effect of set-size (*F*(1,20) = 0.06, *p* = .81). The cluster-based *F*-test identified a significant cluster spanning 1400–2060 ms across widespread electrodes (25 channels: Fp1, AF3, F3, FC1, C3, P7, P3, Pz, O1, Oz, O2, PO4, P4, P8, CP6, C4, T8, FC6, FC2, F4, F8, AF4, Fp2, Fz, Cz; $F_{mass}$ = ${1.45\times10}^{4}$, *p* < .05, $\eta_{mean}= .49$). Yet, follow-up pairwise cluster-based *t*-tests yield no significant effects (all *p* > .05). Taken together, these results suggest that ERP responses during maintenance did not systematically scale with WM load.

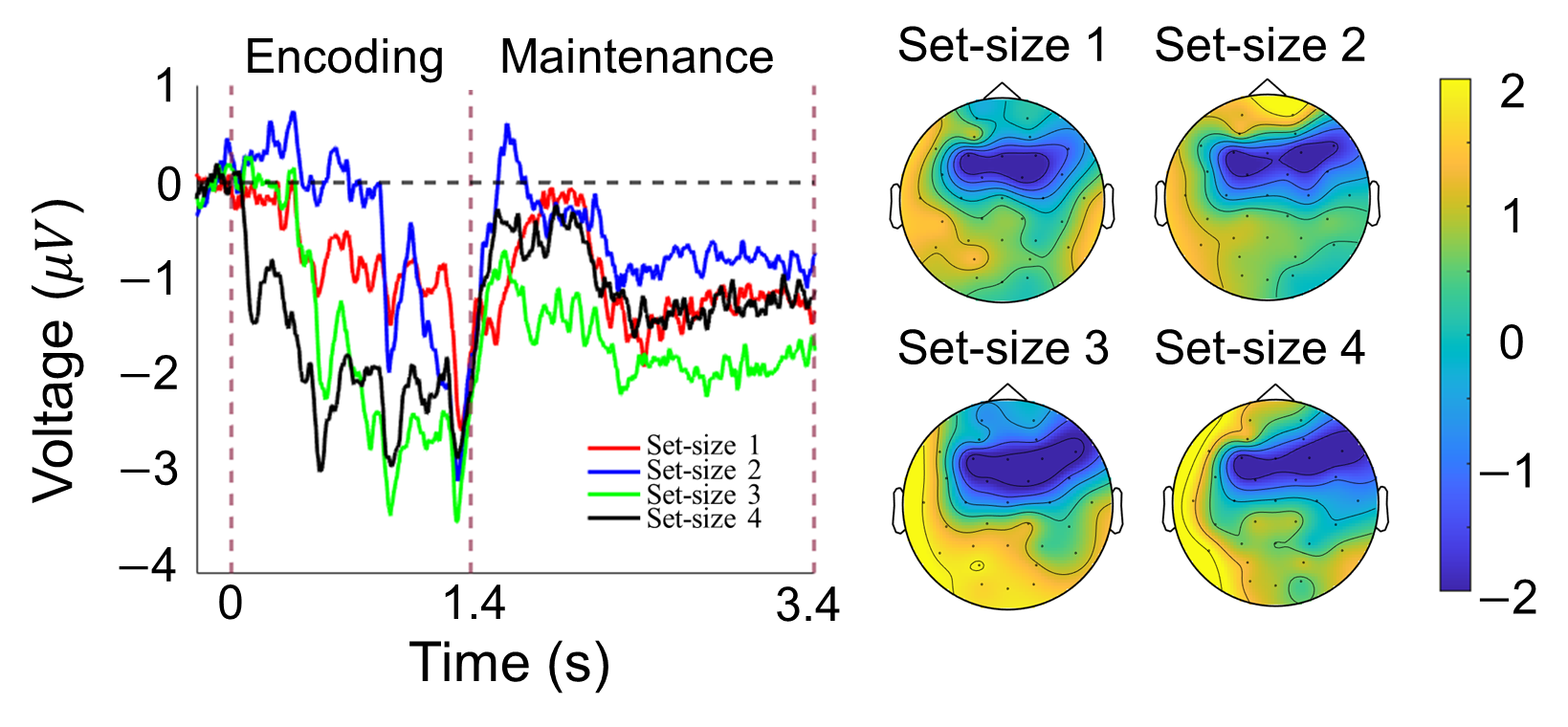

Figure S4. Non-lateralized responses, and their topographical distributions (averaged across the delay), separately for the set-size 1, 2, 3, and 4 conditions. Left: Non-lateralized responses in the time domain data measured in the set-size 1 (red), 2 (blue), 3 (green), and 4 (black) conditions. The vertical dashed lines indicate (from left to right) the onset of the tone sequence, the onset of the maintenance period, and the end of the maintenance period. Right: The four topographical maps depict the magnitude of the non-lateralized responses in set-size 1, 2, 3, and 4 conditions during the maintenance period.

***Multivariate ERP voltage decoding***

To further test whether WM load could be distinguished from scalp voltage patterns, we performed timepoint-by-timepoint multivariate decoding across all electrodes, following the same method reported in the main manuscript for alpha decoding. Cluster-based permutation tests revealed that, overall, set-size could be reliably decoded during the encoding period, from sequence onset until ~ 1600 ms (1 cluster, *p* < .001, $d_{mean}= .80$), but not during the maintenance interval (Figure S5).

We then examined for each pair of set-size conditions individually, whether they evoked discriminable scalp patterns of ERP voltages (i.e., 1 vs 2, 1 vs 3, 1 vs 4, 2 vs 3, 2 vs 4, and 3 vs 4). As shown in the six smaller panels of Figure S5, no significant decoding emerged during the maintenance period. During the encoding phase, we again observed significant decoding for all six set-size comparisons (see Figure S5, small panels), thus further substantiating the quality of the data.

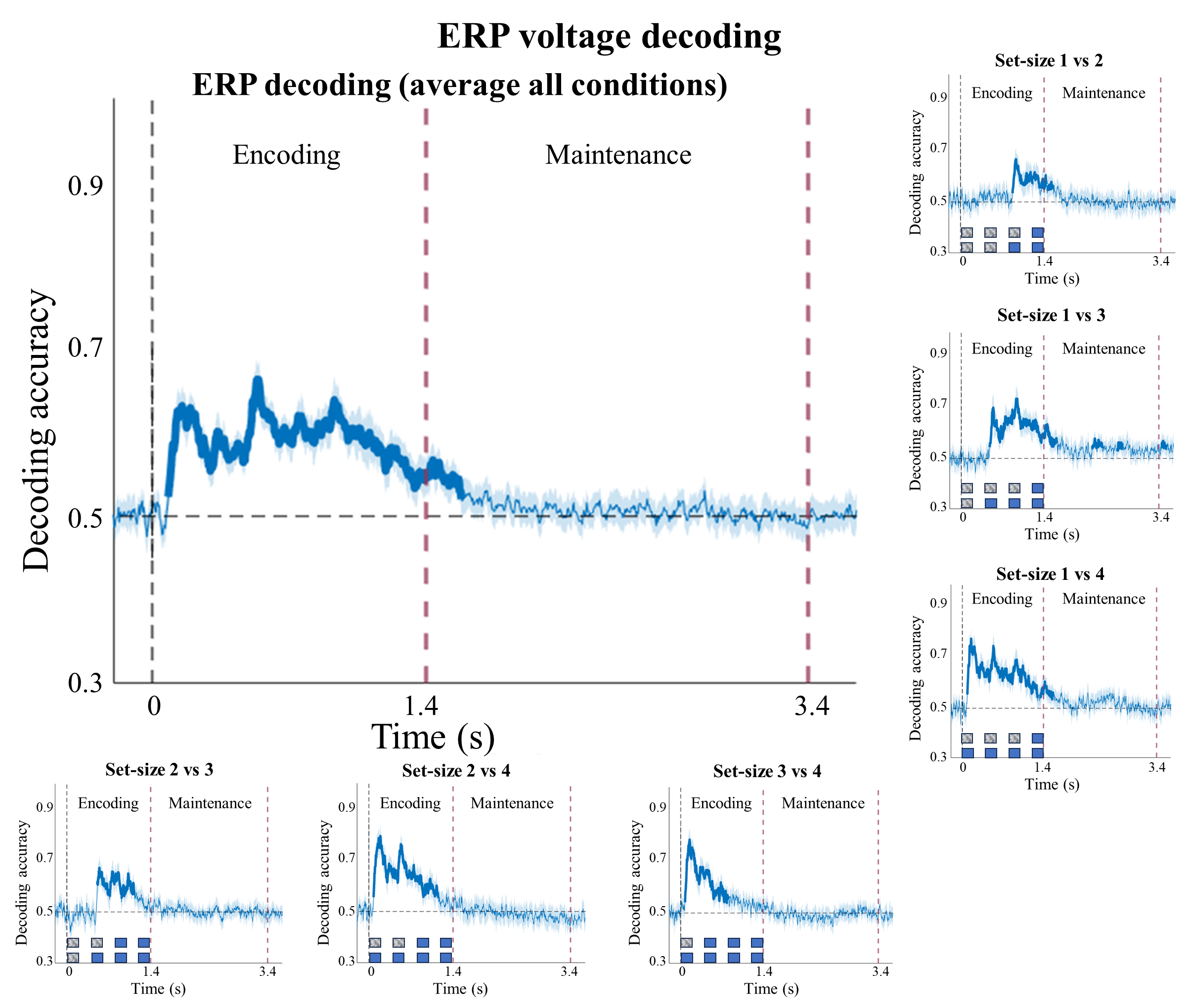

Figure S5. Timepoint-by-timepoint decoding of scalp voltage patterns. All panels depict decoding accuracy (y-axis) as a function of time (x-axis). The big panel depicts decoding accuracy averaged across all six pairwise comparisons (i.e., main effect of set-size), which are shown individually in the surrounding smaller panels. Bold blue lines indicate significant above chance (50%) decoding, based on cluster-based permutation tests to account for multiple comparisons. The vertical dashed-purple lines split time into the encoding and maintenance periods. In the small panels, the blue squares indicate the target tones, while the blurred squares represent the white noise.

Overall, both univariate ERP responses and multivariate decoding results show converging results, suggesting that non-lateralized univariate responses (ERPs) do not scale with set-size during the maintenance period.
